# A bipartite element with allele-specific functions safeguards DNA methylation imprints at the *Dlk1-Dio3* locus

**DOI:** 10.1101/2020.05.22.103796

**Authors:** BE Aronson, L Scourzic, V Shah, E Swanzey, A Kloetgen, A Polyzos, A Sinha, A Azziz, I Caspi, J Li, B Pelham-Webb, H Wichterle, A Tsirigos, M Stadtfeld, E Apostolou

## Abstract

Dysregulation of imprinted gene loci also referred to as loss of imprinting (LOI) can result in severe developmental defects and other diseases, but the molecular mechanisms that ensure imprint stability remain incompletely understood. Here, we dissect the functional components of the imprinting control region of the essential *Dlk1-Dio3* locus (called IG-DMR) and the mechanism by which they ensure imprinting maintenance. Using pluripotent stem cells carrying an allele-specific reporter system, we demonstrate that the IG-DMR consists of two antagonistic regulatory elements: a paternally methylated CpG-island that prevents the activity of Tet dioxygenases and a maternally unmethylated regulatory element, which serves as a non-canonical enhancer and maintains expression of the maternal *Gtl2* lncRNA by precluding *de novo* DNA methyltransferase function. Targeted genetic or epigenetic editing of these elements leads to LOI with either bi-paternal or bi-maternal expression patterns and respective allelic changes in DNA methylation and 3D chromatin topology of the entire *Dlk1-Dio3* locus. Although the targeted repression of either IG-DMR or *Gtl2* promoter is sufficient to cause LOI, the stability of LOI phenotype depends on the IG-DMR status, suggesting a functional hierarchy. These findings establish the IG-DMR as a novel type of bipartite control element and provide mechanistic insights into the control of *Dlk1-Dio3* imprinting by allele-specific restriction of the DNA (de)methylation machinery.

**HIGHLIGHTS:**

- The IG-DMR is a bipartite element with distinct allele-specific functions
- A non-canonical enhancer within the IG-DMR prevents DNA methyltransferase activity
- Targeted epigenome editing allows induction of specific imprinting phenotypes
- CRISPRi reveals a functional hierarchy between DMRs that dictates imprint stability

## INTRODUCTION

More than 100 mammalian genes, most of them found within coregulated clusters, are expressed in a monoallelic, parent-of-origin specific manner (Tucci et al., 2019). This phenomenon, referred to as genomic imprinting, is essential for mammalian development. Imprinted expression is controlled by DNA methylation marks that are established in germ cells in a sex-specific manner at cis-regulatory differentially methylated regions (DMRs) called Imprinting Control Regions (ICR). DMRs within imprinted gene loci are subsequently acted upon by specific transcription factors (TF) and chromatin modifiers to ultimately establish patterns of mono-allelic gene expression (Ferguson-Smith and Bourc’his, 2018). These epigenetic patterns are generally preserved in somatic cells throughout development and in adult tissues and their dysregulation by loss-of-imprinting (LOI) can lead to fetal death or developmental abnormalities as well as other disorders, such as cancer (Kalish et al., 2014). The mechanisms by which ICRs ensure retention of parent-of-origin DNA methylation, known as maintenance of imprinting (MOI), remain incompletely understood.

The *Dlk1-Dio3* locus is a paradigmatic imprinted gene cluster that encodes multiple non-coding and coding transcripts within a region that spans almost one megabase of mouse chromosome 12 (da Rocha et al., 2008). LOI at *Dlk1-Dio3* is associated with severe developmental defects and aggressive malignancies (da Rocha et al., 2008, Jelinic and Shaw, 2007, Khoury et al., 2010, Manodoro et al., 2014) and can also occur during somatic cell reprogramming, resulting in induced pluripotent stem cells (iPSCs) with diminished developmental potential (da Rocha et al., 2008, Carey et al., 2011, Liu et al., 2010, Stadtfeld et al., 2012, Mo et al., 2015). An intergenic DMR or IG-DMR, which resides between the maternally expressed *Gtl2* long non-coding RNA (lncRNA) and the paternally expressed *Dlk1* protein coding gene, has been shown to function as the ICR of *Dlk1-Dio3* (Lin et al., 2003). Similarly to the *H19-Igf2* and *Rasgrf1* ICRs (Kobayashi et al., 2006, Yoon et al., 2005) the IG-DMR is methylated on the paternal allele and appears to control methylation of a secondary DMR that spans the promoter and exon 1/intron 1 of *Gtl2* (“Gtl2 DMR”) (Figure 1A).

**Figure 1:**
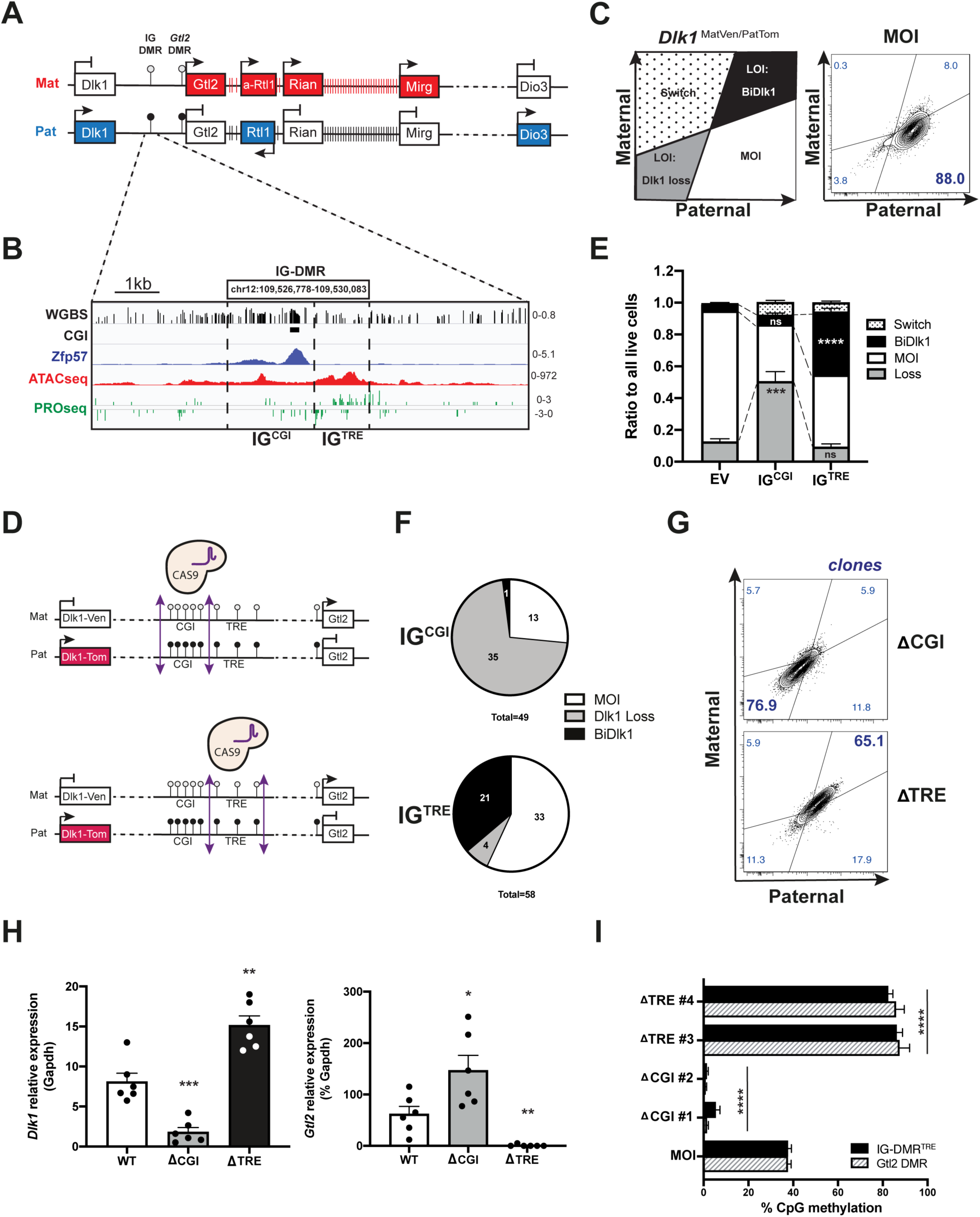
a bipartite Intergenic DMR element regulates genomic imprinting at *Dlk1-Dio3*. (**A**) Schematic representation of the *Dlk1-Dio3* murine locus highlighting maternally (red) and paternally (blue) expressed genes (not in scale). DMR=Differentially Methylated Region, IG=intergenic, Mat=maternal, Pat=paternal. Lollipops represent the DNA methylation state: dark= methylated and white= unmethylated. (**B**) The first 2kb of IG-DMR is characterized by a high CpG density (WGBS) including a canonical CpG island (IG^CGI^) which colocalizes with ZFP57 DNA binding. On the other hand, the remaining distal part (IG^TRE^) constitutes a highly accessible (ATAC-seq) and actively transcribed region (PRO-seq) (see also Figure S1A). IG-DMR coordinates (mm10) are boxed. (**C**) Schematic representation of the distinct phenotypic populations that can be detected by flow cytometry using a *Dlk1* iPSC reporter, where paternal *Dlk1* is linked to tdTomato and maternal *Dlk1* to mVenus. A phenotypic example of normally imprinted (MOI) sample is provided (right). (**D**) Schematic representation of IG^CGI^ and IG^TRE^ deletions performed by CRISPR/Cas9 technology. (**E**) Relative ratios of cell populations with specific Dlk1 expression patterns, as represented in (C), upon targeting of either IG^CGI^ or IG^Enh^. Empty vector (EV) was used as negative control. Biological replicates used: N=8 for EV, N=8 for IG^CGI^ and N=6 for IG^TRE^. Statistical significance of differences for each population relative to the EV (dashed lines) was calculated using a two-tailed unpaired Student’s *t*-test. (**F**) Pie charts showing the ratios of the different phenotypes observed by FACS analysis of individual clones isolated after CRISPR-Cas9 targeting of IG^CGI^ and IG^Enh^. (**G**) Representative FACS plots showing the LOI-Dlk1loss and the LOI-BiDlk1 phenotypes of individual clones with confirmed deletion of IG^CGI^ (ΔCGI) and IG^TRE^ (ΔTRE) respectively. (**H**) RT-qPCR analysis showing *Dlk1* and *Gtl2* expression relative to *Gapdh* in Retinoic Acid (RA)-differentiated clones (N=6) with the respective genotypes (WT: wild type, ΔCGI and ΔTRE. Statistical difference relative to WT was calculated using a two-tailed unpaired Student’s *t*-test. (**I**) Percentage of DNA methylation at the IG-DMR and the *Gtl2* DMR as assessed by pyrosequencing at individual CpG resolution. N=2 clones per genotype are represented. Two-tailed Paired Student’s *t*-test was used to calculate significance relative to MOI levels. ***For all panels:*** Asterisks indicate significance: * p□≤□0.05, ** p□≤□0.01, *** p□≤□0.001, **** p□≤□0.0001. n.s: not significant. Error bars represent +/-SEM.

*Gtl2* has been shown to repress the maternal *Dlk1* gene in *cis* through recruitment of the Polycomb Repressive Complex II (PRC2) (Zhao et al., 2010, Das et al., 2015, Kaneko et al., 2014, Sanli et al., 2018). This suggests that the control of *Gtl2* expression by the IG-DMR (Lin et al., 2003, Kota et al., 2014, Luo et al., 2016, Das et al., 2015) is essential for maintenance of imprinting at *Dlk1-Dio3*. However, how the IG-DMR achieves this regulation in an allele-specific manner elusive. Targeted deletions of the IG-DMR (∼4kb region) in mice have shown that transmission of maternal deletion results in LOI, and specifically in paternalization of the maternal allele, including loss of *Gtl2* and bi-allelic expression of *Dlk1* (Lin et al., 2003). However, transmission of the paternal deletion results in no phenotype (Lin et al., 2003, Das et al., 2015), which is surprising given that the paternal IG-DMR becomes “imprinted” by DNA methylation in primordial germ cells (PGCs) (Sato et al., 2011, Nowak et al., 2011, SanMiguel and Bartolomei, 2018). Different phenotypes were recently reported upon deletion of a 216 bp tandem repeat CpG Island (CGI) within the IG-DMR, where the repressive zinc finger protein Zfp57 binds. Paternally transmitted deletion of this element resulted in maternalization of the paternal allele, whereas maternal transmission of the CGI deletion had no phenotype (Saito et al., 2018, Hara et al., 2019).

These apparently contradictory findings suggested to us that the IG-DMR might be a complex genomic element with multiple cis-regulatory regions that coordinate allele-specific gene expression. We therefore decided to dissect the molecular regulatory logic of imprint maintenance at *Dlk1-Dio3* in pluripotent stem cells (PSCs), a cell type that represents a tractable model system to investigate epigenetic mechanisms including imprinting (Swanzey et al., 2020). Our results show that the IG-DMR is a bipartite element that maintains imprinting by stabilizing the germ line-specific DNA methylation state at *Dlk1-Dio3*. This is achieved by the allele-specific function of two antagonistic cis-regulatory regions within the IG-DMR, which restrict the activity of Dnmt3s and Tet proteins and thereby operate as activator or repressor of the *Gtl2* DMR, respectively. Allele-specific modulations of these elements was sufficient to induce specific and opposing expression phenotypes and epigenotypes, both indicative of LOI. Intriguingly, we observed that epigenetic repression of the *Gtl2* promoter caused bi-allelic *Dlk1* expression without affecting methylation at the IG-DMR. However, this LOI phenotype was unstable and reverted to MOI over time. Therefore, in addition to resolving the complex composition of the IG-DMR, our findings reveal new regulatory principles of the functional hierarchy between genomic elements operational at a gene cluster essential for mammalian development.

## RESULTS

### The IG-DMR is a bipartite element with two distinct functions

To gain insights into the regulatory mechanisms underlying the maintenance of *Dlk1-Dio3* imprinting (Figure 1A), we first assembled in-house (Liu et al., 2017, Di Giammartino et al., 2019) and published datasets (Williams et al., 2011, Shi et al., 2019), including datasets from the CODEX database (Sanchez-Castillo et al., 2015), of DNA methylation, chromatin accessibility, nascent RNAs, histone modifications and TF binding profiles at the IG-DMR in mouse ESCs (Takada et al., 2002, Kobayashi et al., 2006, Hiura et al., 2007). This revealed a pronounced dichotomy with respect to the position and nature of epigenomic features within the IG-DMR (chr12: 109,526,778-109,530,083 mm10) (Figures 1A-B and S1A). The 2kb region closer to the *Dlk1* gene exhibits a high CpG density including a CpG island (CGI) with conserved tandem repeats between human, mouse and sheep (Paulsen et al., 2001), binding of the repressive KRAB domain containing zinc-finger protein Zfp57 (Quenneville et al., 2011, Luo et al., 2016, Riso et al., 2016, Shi et al., 2019) (Figures 1A-B) and components of chromatin-modifying complexes such as PRC2 (Figure S1A). In contrast, the 1kb of the IG-DMR closest to the *Glt2* gene is associated with an open chromatin state enriched for the activating H3K27ac mark, nascent bidirectional transcription, the binding of multiple pluripotency-associated TFs and general transcription regulators, consistent with a Transcriptional Regulatory Element (TRE) function (Danko et al., 2015) (Figure S1A). These data suggest that the IG-DMR may consist of two distinct regulatory elements, which we will refer to as IG^CGI^ and IG^TRE^.

To functionally interrogate potentially distinct roles of the IG^CGI^ and IG^TRE^ in regulating *Dlk1-Dio3*, we deleted each of these elements in iPSCs harboring a previously described allele-specific *Dlk1* reporter system (Swanzey and Stadtfeld, 2016). This transgenic system, in which maternal and paternal *Dlk1* alleles are transcriptionally linked to distinct fluorescent reporters (mVenus and tdTomato respectively) (Figure 1C), has been shown to faithfully capture the imprinting status of the entire *Dlk1-Dio3* locus (Swanzey and Stadtfeld, 2016). Since undifferentiated iPSCs do not express appreciable levels of *Dlk1*, we optimized our flow cytometry analysis approach by combining retinoic acid (RA) induced differentiation with antibody staining against the neuroectodermal marker CD24 and gating on CD24^+^ cells (Semrau et al., 2017) (Figure S1B). This strategy allowed us to reliably identify instances of LOI that lead either to bi-allelic expression (LOI-BiDlk1) or complete silencing (LOI-Dlk1 Loss) of *Dlk1*. Reporter iPSCs were transiently transfected with plasmids expressing Cas9 and pairs of guide RNAs (gRNAs) targeting the 5’- and 3’-ends of either IG^CGI^ and IG^TRE^ (Figure 1D, Table S1 and S3). Strikingly, while targeting of the IG^CGI^ resulted in an increased proportion of cells that lost *Dlk1* expression (LOI-Dlk1Loss), targeting of the IG^TRE^ led to bi-allelic expression of *Dlk1* (LOI-BiDlk1) (Figure 1E). We confirmed that the observed phenotypes were induced by specific deletion of the respective elements, using clonal lines obtained by single cell sorting of respective bulk populations (Figure 1F). This established LOI-Dlk1 Loss as the predominant LOI phenotype in IG^CGI^ targeted clones and LOI-BiDlk1 in IG^TRE^ targeted clones (Figure 1G). Genotyping confirmed mono- or bi-allelic deletion of the targeted regions, while clones with MOI had unedited genomes (Figure S1C and Table S2). These phenotypes were further validated in the context of directed differentiation towards motor neurons (Wichterle et al., 2002, Novitch et al., 2001) (Figures S1D,E). Quantitative PCR confirmed the *Dlk1* expression changes observed with flow cytometry and showed an inverse expression pattern between *Dlk1* and the maternally expressed lncRNA *Gtl2* (Figure 1H and Table S2). Bisulfite sequencing of ΔIG^CGI^ and ΔIG^TRE^ clones showed that both the IG-DMR itself and the *Gtl2 DMR* had completely lost methylation upon deletion of IG^CGI^, while both control elements were fully methylated upon elimination of IG^TRE^ (Figure 1I). Taken together, these data show that the IG-DMR consists of two distinct regulatory elements with essential – but antagonistic – roles in imprinting maintenance. Deletion of these elements is sufficient to induce pronounced and specific epigenetic and transcriptional changes throughout the locus and induce either bi-maternal or bi-paternal LOI.

### IG^CGI^ and IG^TRE^ are allele-specific regulators of *Gtl2* methylation and expression

Next, we sought to establish the allelic specificity of IG^CGI^ and IG^TRE^. To that end, we generated iPSCs carrying the aforementioned allele specific *Dlk1* reporter system in a hybrid JF1/B6 F1 background (Figure 2A and Table S3). The JF1 strain carries several SNPs in the IG-DMR compared to the B6 genetic background (Koide et al., 1998) and ESCs from this strain maintain normal imprinting upon culture (Lee et al., 2018). By using gRNAs designed to target sequences with strain-specific SNPs within the first 3 base-pairs downstream of the PAM sequence, we performed allele-specific CRISPR-Cas9 deletion of the IG^CGI^ and IG^TRE^ regions (Figure S2A and Table S1, S3). Derivation and analysis of individual clones by flow cytometry demonstrated that deletion of the paternal IG^CGI^ (Pat^CGI^) was responsible for the Dlk1 loss phenotype, while the BiDlk1 phenotype was induced by maternal IG^TRE^ (Mat^TRE^) deletion (Figures 2B, S2B). Genotyping confirmed allele-specific deletions (Figures S2C-F and Table S2). While deletion of Mat^CGI^ did not result in a phenotype, some clones (10 out of 56) from the Pat^TRE^ targeting did result in the same BiDlk1 phenotype observed upon targeting the maternal allele. However, investigation of these clones revealed indels on the maternal allele (data not shown) indicating allelic promiscuity of Cas9 (Capon et al., 2017), supporting that deletion of the Pat^TRE^ does not result in LOI.

**Figure 2:**
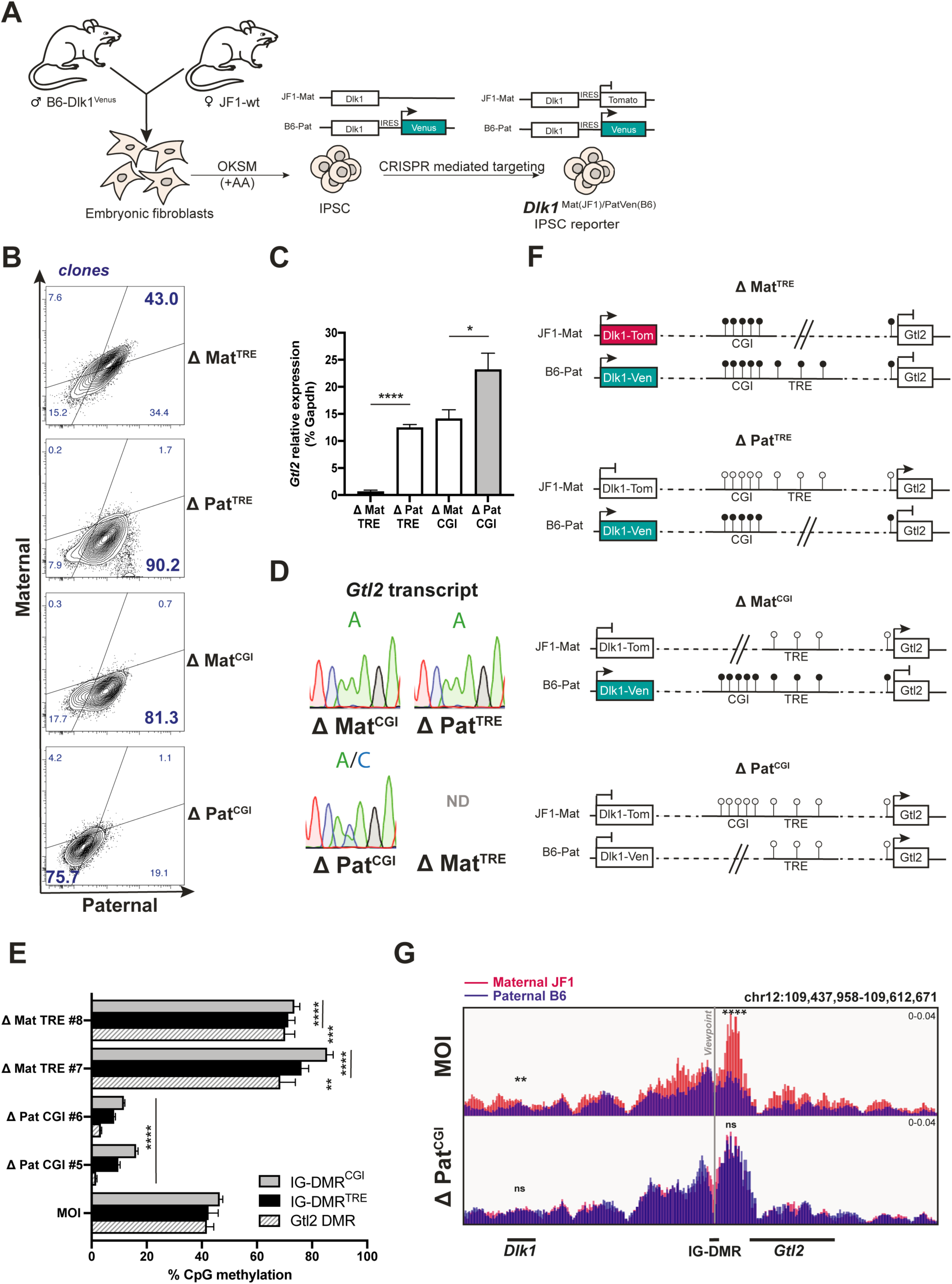
Dual role of IG-DMR in allele specific regulation of *Dlk1-Dio3* locus. (**A**) Schematic illustration of the CRISPR/Cas9 mediated targeting of the maternal *Dlk1* 3’UTR with an IRES-tdTomato cassette. B6= *Mus musculus* C57Bl/6 and JF1=*Mus molossinus* mouse strains. OKSM=Oct4, Klf4, Sox2 and Myc reprogramming factors. AA= Ascorbic Acid. (**B**) Representative FACS plots of clones carrying allele-specific CRISPR/Cas9 deletions of the IG^CGI^ and IG^TRE^. (**C**) RT-qPCR showing the allele-specific *Gtl2* expression levels relative to *Gapdh* (N=2 clones). Statistics was calculated using a two-tailed unpaired Student’s *t*-test. (**D**) A heterozygous SNP in the *Gtl2* RT-qPCR amplicon allows for evaluation of allele-specific expression of *Gtl2* in representative clones. (**E**) Average percentage of DNA methylation at the IG-DMR and the *Gtl2* DMR as assessed by pyrosequencing at individual CpG resolution. N=2 clones with maternal deletion of IG^TRE^ and N=2 clones with paternal deletion of IG^CGI^ are assessed. Two-tailed Paired Student’s *t*-test was used to calculate significance relative to MOI levels. (**F**) Schematic abstract of the molecular changes (DNA methylation and expression) at the Dlk1-Gtl2 locus upon allele-specific CRISPR/Cas9 deletions of the IG^CGI^ and IG^TRE^. (**G**) Allele-specific 4C-seq, using the IG-DMR as a viewpoint, demonstrates maternal interactions in MOI iPSCs and biallelic interactions in Pat^CGI^ clones. Statistics for IG-DMR interaction (Mat vs Pat) with *Gtl2* and *Dlk1* are shown at chr12:109540996-109568650 (FDR= 2.3E^−64^ for MOI and 0.760 for Pat^CGI^) and chr12:109453455-109463336 (FDR= 0.015 for MOI and 0.760 for Pat^CGI^), respectively. ***For all panels:*** Asterisks indicate significance: * p□≤□0.05, ** p□≤□0.01, *** p□≤□0.001, **** p□≤□0.0001. n.s: not significant. Error bars represent +/-SEM.

Quantitative PCR analysis and subsequent sequencing of clonal amplicons confirmed the expected reciprocal effects on Pat^CGI^ and Mat^TRE^ deletion on allele-specific *Gtl2* expression while knockout of the allelic counterparts resulted in no changes (Figures 2C-D and Table S2). Consistent with these findings and our results using non-allele specific genetic engineering (Figure 1I), CpGs within the IG-DMR and *Gtl2* DMR were unmethylated upon paternal IG^CGI^ removal (indicating loss of the paternal methylation mark) and fully methylated upon maternal IG^TRE^ removal (indicating methylation of the maternal allele) (Figure 2E). To investigate whether additional chromatin changes occur in the locus, we performed allele-specific chromatin immunoprecipitation (as-ChIP) for the active histone mark H3K27ac followed by Sanger sequencing. While MOI cells strongly enriched for H3K27ac only on the maternal IG-DMR and *Gtl2* DMR, ΔPat^CGI^ clones exhibited bi-allelic H3K27ac and ΔMat^TRE^ clones lost H3K27ac almost entirely (Figure S2G-H and Table S2). Collectively, our results demonstrate that the unmethylated IG^TRE^ maintains *Gtl2* expression and an active chromatin state on the maternal allele, whereas the methylated IG^CGI^ maintains *Gtl2* repression and an inactive chromatin state on the paternal allele (Figure 2F).

We also asked whether the observed epigenetic and transcriptional changes were accompanied by 3D chromatin reorganization around the locus, by using allele-specific 4C-seq. In agreement with recent reports (Lleres et al., 2019), we found that the long-range chromatin contacts around the IG-DMR differ between the maternal and paternal alleles. Specifically, the maternal, unmethylated IG-DMR interacts at high frequency with both *Gtl2* and *Dlk1*, while these interactions are significantly weaker on the paternal allele (Figure 2G). Deletion of the paternal IG^CGI^ perturbed this allele-specific topology and induced a maternal-like conformation on both alleles (Figures 2G and Table S2). On the other hand, deletion of the maternal IG^TRE^ induced a significant overall decrease of interactivity resembling a bi-paternal topology (Figure S2I). These results suggest that DNA methylation of IG^CGI^ might prevent the physical interaction of the IG-DMR with its target genes and that IG^TRE^ is essential for the maintenance of these long-range chromatin contacts. Taken together, our observations show that the paternal IG^CGI^ and maternal IG^TRE^ have distinct regulatory functions that are required to maintain the respective alleles in epigenetic and conformational states consistent with MOI at *Dlk1-Dio3*.

### The bipartite nature of the IG-DMR results in allele-specific suppression of DNA methyltransferase and Tet protein activity

*De novo* DNA methyltransferases and Tet dioxygenases operate at ICRs to establish and erase methylation in germ cells during development (Li and Sasaki, 2011, Bartolomei and Ferguson-Smith, 2011, Plasschaert and Bartolomei, 2014, SanMiguel and Bartolomei, 2018). The allele specific DNA methylation changes that we observed upon IG^TRE^ and IG^CGI^ deletion (Figure 2E) raised the possibility that these enzymes might also be involved in the induction of distinct LOI phenotypes. To test this possibility, we deleted IG^CGI^ or IG^TRE^ in parallel to the gene loci encoding all Tet proteins (*Tet1, Tet2* and *Tet3)* or both Dnmt3 enzymes (*Dnmt3a/b*) in *Dlk1* reporter iPSCs (Figure 3A and Table S1). Subsequent sorting, differentiation and FACS analysis of bulk populations showed a significant amelioration of the LOI-Dlk1 Loss phenotype induced by IG^CGI^ deletion when Tets were simultaneously deleted (Figures 3B-C). Similarly, simultaneous deletion of IG^TRE^ and of *Dnmt3a/b* counteracted the LOI-BiDlk1 phenotype and increased the proportion of iPSCs that retained MOI (Figures 3B-C). These observations suggest that the IG^CGI^ ensures imprinting stability by blocking Tet activity on the paternal allele, while the IG^TRE^ protects imprinting by blocking Dnmt3s activity on the maternal allele. Genotyping of targeted regions and surveyor assays in bulk populations confirmed the respective deletions and indels by CRISPR/Cas9 and showed amplicons of equal strength, demonstrating similar degrees of targeting efficiency across experimental conditions (Figures S3A-B and Table S2). Importantly, ablation of Dnmt3s or of the Tet enzymes without simultaneous removal of either IG^CGI^ or IG^TRE^ showed no effects on imprinting stability within our experimental time window (5-6 passages). This indicates that neither of these enzymatic activities are required for maintenance of imprinting at *Dlk1-Dio3* in the presence of an intact IG-DMR. Rather, the major function of the IG-DMR appears two-fold: (i) IG^TRE^-mediated prevention of *de novo* DNA methylation on the maternal allele and (ii) IG^CGI^-mediated prevention from active DNA demethylation on the paternal allele by Tets.

**Figure 3:**
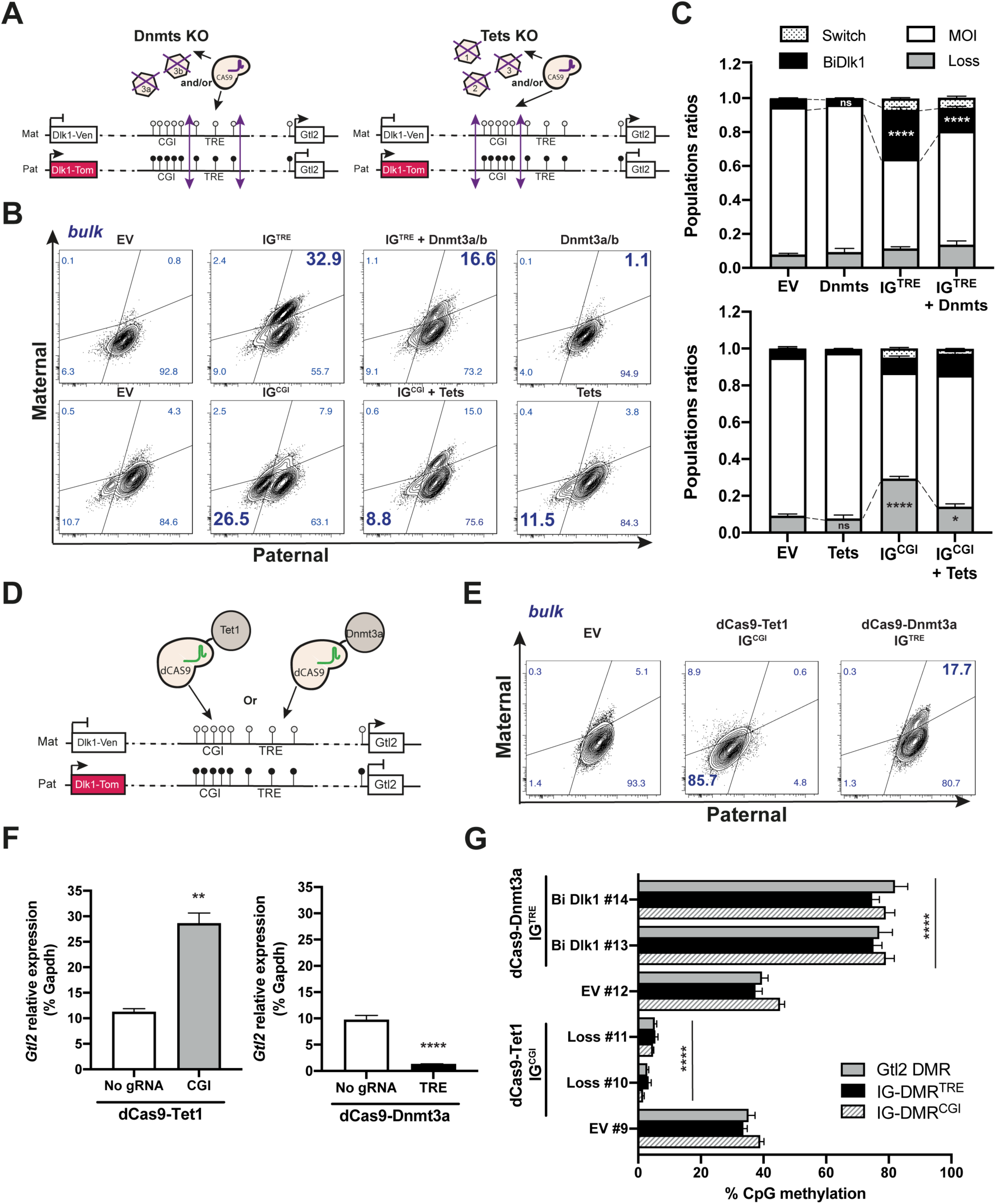
Maintenance of IG-DMR imprinting is mediated by prevention of DNA methyltransferases and Tet proteins function. (**A**) Schematic representation of CRISPR/dCas9 mediated genome targeting of the IG-DMR elements in combination with targeting of Dnmt3a/b genes (Dnmts KO) and Tet1/2/3 genes (Tets KO). (**B**) Representative FACS plots of bulk populations upon each targeting strategy. (**C**) Quantification of phenotypic FACS populations from N=6 targeting experiments. Statistical significance for each population compared to EV (dashed line) was calculated using two-tailed unpaired Student’s *t*-test. (**D**) Schematic illustration of our strategy to target dCas9-Tet1 to the IG^CGI^ or dCas9-Dnmt3a to the IG^TRE^ and (**E**) Representative FACS plots of bulk populations upon each targeting strategy. (**F**) RT-qPCR showing that *Gtl2* RNA expression levels in dCas9-Tet1 and dCas9-Dnmt3a reporter show an inverse correlation with *Dlk1* reporter expression, as expected. Statistical significance was calculated by two-tailed unpaired Student’s *t*-test. (**G**) DNA methylation analysis shows that methylation is lost upon dCas9-Tet1 targeting to the IG^CGI^ while it is gained upon dCas9-Dnmt3a targeting to IG^TRE^. Statistical significance for each DMR was calculated using two-tailed paired Student’s *t*-test relative to the respective EV clone (clone #9 for dCas9-Tet1 targeted IG^CGI^ and clone #12 for dCas9-Dnmt3a targeted IG^TRE^, see also Table S4) ***For all panels:*** Asterisks indicate significance: * p□≤□0.05, ** p□≤□0.01, *** p□≤□0.001, **** p□≤□0.0001. n.s: not significant. Error bars represent +/-SEM.

### Forced gain or loss of DNA methylation at the IG-DMR is sufficient to induce LOI

The results described above demonstrate that the protection of allele-specific DNA methylation states by the IG^CGI^ and IG^TRE^ are necessary to maintain imprinting. We therefore asked whether local DNA methylation changes at these respective elements could phenocopy the effect of specific genetic deletions on imprint stability. We generated *Dlk1* reporter iPSCs that express a catalytically deactivated Cas9 (dCas9) fused to the catalytic domain of either TET1 or DNMT3A (Tables S2 and S3) (Xu et al., 2016, Choudhury et al., 2016, Morita et al., 2016, Verma et al., 2018). We then stably expressed multiplexed gRNAs targeting either three Zfp57 binding sites within the IG^CGI^ or the sites of nascent transcription of the IG^TRE^ (Figure 3D and Table S1). We observed that targeting of dCas9-Tet1 to the IG^CGI^ resulted in conversion of almost all cells to the LOI-Dlk1 Loss phenotype (Figure 3E), recapitulating our observations upon deletion of the Pat^CGI^ (Figure 2B). This phenotype was accompanied by loss of DNA methylation on both IG-DMR and *Gtl2* DMR and by *Gtl2* upregulation (Figures 3F, G and Table S2). On the other hand, targeting of dCas9-Dnmt3a to the IG^TRE^ induced a LOI-BiDlk1 population with respective hypermethylation and loss of *Gtl2* expression (Figures 3E-G). Taken together, these experiments demonstrate that local changes in DNA methylation at the IG^CGI^ and IG^TRE^ are sufficient to affect the methylation status at the distal *Gtl2* DMR and perturb imprinted gene expression. This supports the notion that protection against inadvertent Dnmt3/Tet enzymatic activity is a major function of the subregions of the IG-DMR.

### DMR-specific targeting of CRISPRi results in either transient or irreversible LOI

So far, our results have shown that allele-specific genetic or epigenetic manipulation of the specific elements within the IG-DMR induces LOI phenotypes that correlate with methylation and transcriptional changes at the *Gtl2*. To test whether the regulatory function of the IG-DMR can be overridden by direct silencing of *Gtl2*, we generated stable reporter iPSCs expressing the transcriptional repressor dCas9-BFP-KRAB (CRISPRi) and gRNAs that target either the IG^CGI^, the IG^TRE^ or the *Gtl2* DMR (Figure 4A and Table S1). Consistent with dCas9-mediated epigenetic editing of the IG-DMR (Figures 3D-G), targeting of CRISPRi to either the IG^CGI^ or IG^TRE^ resulted in hypermethylation of the IG-DMR and *Gtl2* DMR as well as *Gtl2* repression (Figures 4B-D and S4A, B). Targeting of the *Gtl2* DMR similarly triggered a high degree of LOI-BiDlk1, but in contrast to IG-DMR targeting, only induced local *Gtl2* DMR hypermethylation without affecting the methylation status of the IG-DMR (Figure 4D). This demonstrates that targeted repression of *Gtl2* is sufficient to activate maternal *Dlk1* without acquisition of DNA methylation at the IG-DMR. To test whether the absence of IG-DMR hypermethylation might affect the stability of this LOI phenotype, we made use of the observation that our lentiviral CRISPRi, without continuous resorting or reselection, is silenced over time in PSCs. We established BFP^+^ and BFP^−^ subclones from originally BFP^+^ (i.e. dCas9-BFP-KRAB expressing) cells, followed by extensive passaging allowing for potential reversal of epigenetic effects after dCas9-KRAB (Mandegar et al., 2016). This was done in cells where CRISPRi was targeted either to the IG^CGI^ or to the *Gtl2* DMR. FACS analysis showed that all CGI-targeted clones retained their LOI-BiDlk1 phenotype and hypermethylation of both IG-DMR and *Gtl2* DMR, independently of their BFP expression status (Figures 4E-F). This demonstrates that the change in imprinting status was irreversible. In contrast, three out of six *Gtl2*-targeted BFP^−^ clones partially reverted to MOI (Figures 4E-F and S4C). This partial rescue was not seen in clones that retained CRISPRi expression (BFP^+^). Bisulfite sequencing of the MOI population of partially rescued clones showed reestablishment of normal methylation levels (∼50%) at the *Gtl2* promoter. Interestingly, the remaining BiDlk1 population of rescued clones showed intermediate levels of methylation that were ∼10% reduced compared to clones still expressing CRISPRi, suggesting an ongoing DNA demethylation that could reach MOI levels upon further passaging (Figure 4G). Furthermore, qPCR analysis confirmed that *Gtl2* expression was significantly increased in rescued *Gtl2-*targeted clones compared to CGI-targeted clones that had lost CRISPRi expression (Figure 4H and Table S2). These results suggest that although modulating the activity or methylation of the *Gtl2* promoter can transiently alter gene expression at *Dlk1-Dio3*, the DNA methylation status of the IG-DMR is the determining factor for long-term imprinting stability of this cluster.

**Figure 4:**
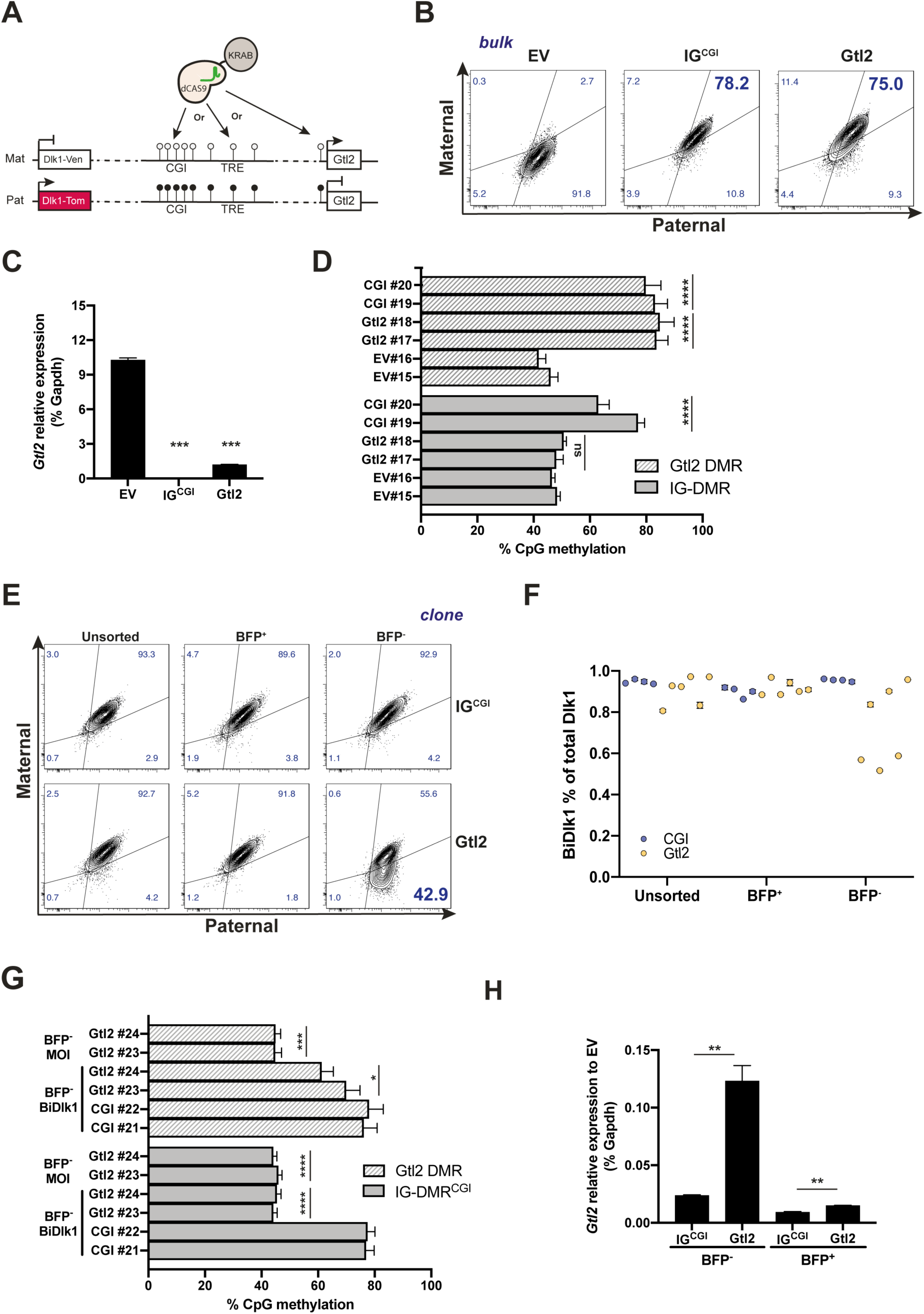
IG-DMR methylation status dictate *Dlk1-Dio3* imprinting stability. (**A**) Strategy to target dCas9-BFP-KRAB to either IG^CGI^ or IG^TRE^ or to the *Gtl2* DMR. (**B**) Representative FACS plots of bulk populations, showing that targeting of either element results in LOI-BiDlk1 population (see also Figure S4A). (**C**) RT-qPCR analysis showing downregulation of *Gtl2* expression in the targeted cells compared to EV. Statistical significance was calculated by two-tailed unpaired Student’s *t*-test. (**D**) Average percentage of methylated CpGs at the IG-DMR and Gtl2 DMR as determined by bisulfite sequencing analysis. Note that targeting dCas9-BFP-KRAB to the *Gtl2* DMR does not result in hypermethylation at the IG-DMR. Statistical analysis of each DMR (and each clone) is compared to the EV #15 and 16 using two-tailed paired Student’s *t*-test. (**E**) Example FACS plots from a dCas9-BFP-KRAB CGI-targeted clone and a rescued *Gtl2*-targeted clone at passage 15 (p15). The *Gtl2*-targeted clone shows that approximately half of the cells have reset MOI. (**F**) Quantificantion of FACs results showing the percentage of LOI-BiDlk1 population in N=4 dCas9-BFP-KRAB CGI-targeted clones and N=6 *Gtl2*-targeted clones at p15. (**G**) Bisulfite sequencing analysis of MOI and BiDlk1 populations of two rescued dCas9-BFP-KRAB targeted clones shows that normal DNA methylation levels (∼50%) at the IG-DMR were reestablished in the MOI population. Statistical significance was calculated by two-tailed paired Student’s *t*-test using CGI clones #21 and #22 as controls (see Table S4). (**H**) RT-qPCR analysis showing that the BFP^−^ population of *Gtl2*-targeted clones shows a drastic upregulation of *Gtl2* compared to IG^CGI^ BFP-. Statistical analysis of Gtl2 and IG^CGI^ are compared within each BFP^−^ or BFP^+^ population using two-tailed paired Student’s *t*-test. ***For all panels:*** Asterisks indicate significance: * p□≤□0.05, ** p□≤□0.01, *** p□≤□0.001, **** p□≤□0.0001. n.s: not significant. Error bars represent +/-SEM.

## DISCUSSION

In this study, we combine genetic engineering with targeted epigenetic editing in mouse pluripotent stem cells to dissect the regulatory mechanisms that protect imprint stability at *Dlk1-Dio3*, a gene cluster essential for mammalian development. Our findings extend prior studies that have identified the IG-DMR as the critical control element of *Dlk1-Dio3* (Lin et al., 2003, Nowak et al., 2011, Kota et al., 2014, Luo et al., 2016, Wang et al., 2017, Saito et al., 2018) and we provide unique insights into its structure and mechanisms of action that can be exploited to modulate *Dlk1-Dio3* imprinting status in a predictable fashion.

By allele-specific CRISPR knockout experiments, we revealed that the IG-DMR is a bipartite element, composed of a repressive element (IG^CGI^) that functions on the paternal allele and an activating element (IG^TRE^) on the maternal allele, both of which are necessary to maintain allele-specific gene expression over distance (Figure S4D). Deletion of each element from the respective allele results in either bi-paternal or bi-maternal LOI, while reciprocal deletions show normal imprinting. Furthermore, we show that the two elements of IG-DMR independently safeguard the imprinted DNA methylation state of *Dlk1-Dio3* by restricting the competing activities of Dnmts and Tets on the locus in an allele-specific manner. All these features document that the IG-DMR is distinct from other previously described composite ICRs, which serve as methylation-sensitive insulator boundaries, as in the case of *Rasgrf1* and *Snrpn* (Yoon et al., 2005, Bartolomei, 2009, Rabinovitz et al., 2012, Hsiao et al., 2019).

It has previously been suggested that the IG-DMR functions as an enhancer for *Gtl2* expression (Lin et al., 2003, Kota et al., 2014, Das et al., 2015, Luo et al., 2016, Wang et al., 2017). Consistent with these studies, we found that deletion of the maternal unmethylated IG^TRE^, or of the entire IG-DMR (data not shown), resulted in hypermethylation of the *Gtl2* DMR, silencing of *Gtl2* and LOI-BiDlk1. However, based on our functional experiments and the unique genomics features of this bipartite unit, we argue that IG-DMR operates as a non-canonical enhancer. Indeed, simultaneous deletion of the IG^TRE^ and loss-of-function mutation of Dnmt3a/b led to the surprising finding that the IG^TRE^ is dispensable for MOI in the absence of *de novo* methyltransferase activity. This result suggests that, rather than directly activating *Gtl2*, the maternal IG^TRE^ allows *Gtl2* expression by preventing *de novo* DNA methylation of the *Gtl2* DMR and thereby upholds MOI. Intriguingly, some of the molecular characteristics of the IG-DMR resemble the features of so-called orphan CGI, which are distal to promoters and are characterized by active enhancer marks, such as H3K27ac, topological interactions with nearby target genes and overlapping Zfp57 and Mll1 consensus motifs (Quenneville et al., 2011, Anvar et al., 2016, Bae et al., 2016, Bina, 2017, Bell and Vertino, 2017, Mendizabal and Yi, 2016). Importantly, similarly to IG-DMR, the activity of orphan CGIs is controlled by DNA methylation (Illingworth et al., 2010) and is frequently dysregulated in cancer (Bell and Vertino, 2017). Future experiments will be required to identify functional commonalities between IG-DMR and orphan CGI enhancers and investigate whether such non-canonical enhancer function is a more general mechanism of gene expression control in development and disease.

Our genetic engineering experiments established the ability of IG^CGI^ and IG^TRE^ to prevent *de novo* DNA methylation and demethylation, respectively. Several trans-acting factors could mediate these functions (Figure S4D). A prime candidate for the IG^CGI^ is the KRAB domain containing zinc finger ZFP57, which has been shown to bind to the methylated IG-DMR in the region of the IG^CGI^ (Riso et al., 2016, Luo et al., 2016, Strogantsev et al., 2015, Shi et al., 2019) and maintains closed chromatin by recruiting repressive complexes, such as DNA and H3K9 methyltransferases (Quenneville et al., 2011, Riso et al., 2016). In agreement with the importance of ZFP57 for the paternal IG-DMR, CRISPR/Cas9 mediated knockout of this protein in allele-specific *Dlk1* reporter iPSCs resulted in IG-DMR hypomethylation and loss of *Dlk1* expression, resembling IG^CGI^ deletion (data not shown). Moreover, knockdown of *Zfp57* has been shown to cause accumulation of 5-hydroxymethylcytosine (5hmC) at the IG-DMR (Coluccio et al., 2018), which suggests that this protein might be essential to prevent Tet enzymes activity on the paternal IG-DMR. Simultaneous deletion of ZFP57 and TET enzymes could resolve this question. The factors and mechanisms that prevent hypermethylation of the maternal IG^TRE^ remain elusive. Our CRISPRi experiments support that the active chromatin and transcriptional status of IG^TRE^ is necessary for protecting the maternal allele from *de novo* DNA methylation both locally and on the *Gtl2* DMR. Indeed, active histone marks (Rose and Klose, 2014) as well as binding of transcription factors or the activating complex CBP/p300 have been shown to protect CpG sites from DNMT3A/B activity (Straussman et al., 2009, Gebhard et al., 2010, Lienert et al., 2011) (Zhang et al., 2017). Similarly, the nascent non-coding transcripts of the locus (enhancer RNA and/or Gtl2 non-coding RNA) may also play protective roles, as has been previously suggested (Kota et al., 2014), by either interacting with PRC2 (Das et al., 2015) or Dnmt3a/b and Dnmt1 (Zhao et al., 2016, Morlando and Fatica, 2018). Further investigation will be required to dissect the relative contribution of each of these factors in maintenance of imprinting at *Dlk1-Dio3*.

The observation that using dCas9 to target epigenome modifiers to different candidate regulatory regions of *Dlk1-Dio3* led to distinct molecular changes and imprinting phenotypes underscores the power of this approach to dissect the regulatory logic of complex genetic loci (Hsu et al., 2014). Combined, our epigenetic targeting experiments revealed a hierarchical and unidirectional regulation between IG-DMR and *Gtl2* DMR that is substantiated by the following key observations. First, although any targeted modulation of the IG-DMR (either by dCas9-Tet1, dCas9-Dnmt3a or CRISPRi) resulted in methylation changes on both DMRs, targeting of the *Gtl2* DMR affected only local DNA methylation without any effects on IG-DMR, suggesting a “one-way” communication. In addition, although targeting of the *Gtl2* DMR can cause LOI, normal imprinting can be restored by the IG-DMR, suggesting that this element controls long-term imprinting stability. Dissecting the molecular mechanisms (e.g. physical proximity or spreading) underlying this regulatory hierarchy and their degree of conservation among cell types (Alexander et al., 2019) and species would be interesting areas for future investigation, in particular in the context of imprinting disorders.

In conclusion, our study refined the mechanistic understanding of the regulatory logic that safeguards imprinting stability at *Dlk1-Dio3* and revealed ways to perturb this logic in a rationale and predictable manner. We envision that the molecular principles operational at the IG-DMR may serve as paradigms for epigenetic regulation beyond imprinting.

## ACKNOWLEDGEMENTS

We are grateful to A. Melnick, D. Campigli di Giammartino, R. Chaligne and the members of the Apostolou, Tsirigos and Stadtfeld laboratories for critical reading of the manuscript, to N. Ben Chetrit for advice on and evaluation of microscopy experiments and to D. Huangfu and L. Dow for sharing CRISPR–Cas9 vectors and cloning protocols. B.E.A. is supported by the National Health Institutes, National Cancer Institute (NCI) Grant T32 CA203702, L.S. is supported by the Lady TATA Memorial Fund and the Lymphoma Research Foundation, B.P.W. is supported by the NICHD with a T32 (T32HD060600) and an F30 (F30HD097926), as well as by a Medical Scientist Training Program grant from the NIGMS under award number T32GM007739 to the Weill Cornell/Rockefeller/Sloan Kettering Tri-Institutional MD-PhD Program, M.S. is supported by R01GM111852-01 from NIH, E.A. is supported by the Edward Mallinckrodt Jr. Foundation, the NIH Director’s New Innovator Award (grant no. DP2DA043813) and the Tri-Institutional Stem Cell Initiative by the Starr Foundation.

## AUTHOR CONTRIBUTIONS

Conceptualization B.E.A., L.L.S., M.S. and E.A.; Methodology B.E.A., L.L.S., M.S. and E.A.; Formal analysis A.K. and A.P.P.; Investigation: B.E.A., L.L.S., V.S., E.S., A.S., A.A., I.C., J.L., B.P.W. and L.C.; Writing – original draft B.E.A., L.L.S., M.S. and E.A.; Supervision H.W., M.S. and E.A.

## DATA AVAILABILITY

4C-seq data generated were submitted to GEO under the accession number GSE148315.

## STAR METHODS

### Lead Contact and Materials Availability

Further information and requests for resources and reagents should be directed to and will be fulfilled by the Lead Contact, Dr. Effie Apostolou.

### Experimental models and subject details

#### ESC and iPSC cell culture

All cells were cultured at 37°C with 5% CO_2._ C57BL/6J and C57Bl/6J-JF1 iPSC’s or V6.5 ESC’s were cultured on plates coated with 0.2% gelatin on irradiated mouse embryonic fibroblasts (MEF’s) with ES KO medium supplemented with 10% FBS, 10mg recombinant leukemia inhibitory factor (LIF), 0.1mM beta-mercaptoethanol (Sigma-Aldrich), penicillin/streptomycin, 1mM L-glutamine and 1% nonessential amino acids (all from Invitrogen).

### Generating allele-specific iPSC *Dlk1*-reporter in hybrid maternal JF1/paternal B6 background

B6 mice used to generate JF1/B6 iPSCs were on a mixed background between C57BL/6NJ (Jax 005304) and C57BL/6J (Jax 000664). 4µg total of IRES-tdTomato-Neomycin donor (Swanzey and Stadtfeld, 2016) and CRISPR/Cas9 vectors were transfected into Mat-JF1 X Pat-B6-Venus Tail Tip Fibroblasts-derived IPSCs using Lipofectamine 2000 (Invitrogen #11668019). Transfected cells were selected for 4 days with 1mg/ml Geniticin (Invitrogen #10131-035) and plated on DR4 MEFs. Individual clones were picked, expanded and screened for the proper integration of tdTomato and persistence of GFP sequence by PCR. Sole expression of Pat-Venus was confirmed by flow cytometry and proper monoallelic Gtl2 expression was confirmed by RT-qPCR and Sanger sequencing (see complete list of expression primers in Table S2).

### iPSC’s with CRISPR/Cas9 mediated deletions

sgRNA sequences were cloned into the pSpCas9(BB)-2A-GFP (px458) vector under the U6 promoter by digesting the vector with BbsI (NEB #0539) and ligating annealed sgRNA oligonucleotides into the vector with T4 ligase (NEB, M0202L) under standard conditions. px458-sgRNA’s were transfected using Lipofectamine 2000 (Invitrogen #11668019) into 300k iPSC’s on plates coated with 0.2% gelatin. 800 ng DNA was transfected per gRNA supplemented to 4µg total DNA with scrambled px458 vector. iPSC’s were sorted for GFP 48-72 hours post-transfection and plated on MEF feeder plates. Subsequent clonal populations were obtained by single-cell sorting the bulk on 96-well plates on the FACS Aria II (BD biosciences). Confirmation of deletions was performed by genotyping using a 3-primer strategy with 2 primers flanking and 1 primer inside the region of interest. As all primers have a different distance to the ‘breakpoint’, deletions and inversions could be detected by gel electrophoresis after PCR. For allele-specific genotyping, the primers were positioned on a SNP on the 3’ end and contained 1 mismatch in the first 5 nucleotides upstream of the SNP. Indels, generated by transfecting Cas9 with a single gRNA, were confirmed with T7 surveyor assays as described previously (Guschin et al., 2010). All sgRNA oligonucleotides, genotyping and Surveyor primers are listed in supplemental Table S1 and S2 respectively.

### Cloning of multiple gRNA expression vectors for dCas9 expressing iPSC’s

sgRNA’s were cloned into the pLKO5.sgRNA.FCS.PAC-NEO under the U6 promoter as described above. Next, the U6 promoter, sgRNA and scaffold combinations were PCR amplified with primers containing overhangs with recognition sites of various restriction enzymes (see below). These enzymes were chosen in such a manner that a digested amplicon would have a complementary overhang with the next amplicon. Digestion and subsequent ligation of all guides into the vector thus resulted in simultaneous insertion of multiple gRNA’s that are each under the control of a single U6 promoter.

To insert three guides, the vector was digested with XbaI (NEB, R0145), amplicon-1 was digested with XbaI and MfeI (NEB, R0589), amplicon -2 was digested with EcoRI (NEB, R0101) and SalI (NEB, R0138) and amplicon -3 was digested with XhoI (NEB, R0146) and AvrII (NEB, R0174). To insert four guides, the vector was digested with XbaI, amplicon -1 was digested with XbaI and MfeI, amplicon -2 was digested with EcoRI and BglII (NEB, R0144), amplicon -3 with BamH1 (NEB, R3136) and SalI and amplicon -4 was digested with XhoI and AvrII (Dow et al., 2015). Ligation was performed in T4 ligation buffer (NEB B0202) and incubated with T4 Ligase (5 μl, 400U/μl; NEB M0202L).

### Cloning of dCas9 plasmids

To create a vector for stable dCas9 expression in iPSC’s, we chose the lentiviral pHR-SFFV-dCAS9-BFP-KRAB vector (Gilbert et al., 2013) and performed an EcoRI digestion to remove the SFFV promoter. An Ef1α promoter sequence was PCR amplified from pdCas9-VP64 (Addgene #61425) using primers with EcoRI restriction site overhangs. The amplicon was EcoRI digested and ligated to replace the SFFV promoter.

To create pHR-dCas9-Tet1-BSD, we digested the pHR-Ef1α-dCAS9-BFP-KRAB with SbfI-HF (NEB, R3642) and MluI-HF (NEB, R0198) to remove dCas9-BFP-KRAB. We then PCR-amplified a dCas9-Tet1 sequence from pENTRY-dCas9-Tet1CD (Verma et al., 2018) and an IRES-BSD sequence from pMIGR-IRES-BSD. The purified IRES-BSD product was digested with EcoRI. The fragments were ligated using Gibson assembly (NEB E2611L).

To create pHR-dCas9-Dnmt3a-BSD, we PCR amplified the dCas9-Dnmt3a sequence from pdCas9-Dnmt3a-puro (Vojta et al., 2016) and digested pMIGR-IRES-BSD with EcoRI to produce an IRES-BSD fragment. The fragments were ligated using Gibson assembly.

To create gRNA delivery vectors, sgRNAs were cloned into the pLKO5.sgRNA.FCS.PAC (Heckl et al., 2014) with adjustments made to the scaffold according to (Chen et al., 2013). The original puromycin cassette was removed by BamHI-MluI digestion and a neomycin sequence was PCR amplified from a pHR-Dlk1-IRES-tomato vector and inserted with Gibson assembly. Primers used for cloning have been listed in Table S2. All used and constructed plasmids have been assembled in Table S3.

### Lentiviral production and infection

293T cells were transfected with overexpression constructs along with the packaging vectors VSV-G and Delta8.9 using PEI reagent (PEI MAX, Polyscience, 24765-2). The supernatant was collected after 48 and 72 h, and the virus was concentrated using polyethylglycol (Sigma, P4338). V6.5 cells or iPSC’s were infected in medium containing 5 μg/ml polybrene (Millipore, TR-1003-G), followed by centrifugation at 2,500 r.p.m. for 90 min at 32 °C. Media was changed after 8-12h following spinfection.

### Generation of stable dCas9 expressing cell lines

To generate stable dCas9 and gRNA expressing cell lines, IPSCs were transduced with concentrated lentivirus (as described above). 5x 3000 cells were pre-plated on gelatin 0.2% on 5 wells of 96-well plates and transduced each with 10ul concentrated virus in 100ul KO-DMEM with polybrene 1mg/ml. The cells were passaged and selected with 10 μg/ml Blasticidin (Life Tech, A11139-03) (dCas9/Dnmt3a and dCas9-tet1) for 4 days or sorted on the FACS Aria II (dCas9-KRAB-BFP). The selected populations were then transduced again with the pLKO5 lentivirus, containing sgRNA’s against the target of interest, and selected with 1mg/ml geneticin (Invitrogen 10131-035) for 5 days.

### Method details

#### Retinoic acid differentiation assays

For differentiation assays, 10K cells/cm2 were plated in standard DMEM supplemented with 10% FBS, 0.1mM beta-mercaptoethanol (Sigma-Aldrich), penicillin/streptomycin, 1mM L-glutamine and 1% nonessential amino acids (all from Invitrogen). 0.4 μg/ml retinoic acid (Sigma-Aldrich, R2625) was added followed by daily media changes for six days.

#### Flow cytometry

To assess the proportion of mVenus and tdTomato in the established reporter cell lines, a single-cell suspension was filtered, stained with anti-mouse CD24a/APC-eFluo780 antibody (Affymetrix 47-0242-82) and assessed on the BD Aria or FACS Canto II. Analysis was done in FlowJo (BD Biosciences).

#### qPCR

Total mRNA was extracted from pre-plated iPSC’s and RA differentiated cells using Qiagen RNeasy kit (Qiagen 74106). On-column DNA digestion was performed on all samples with RNAse free DNAse set (Qiagen 79254). RNA was reverse transcribed with random hexamers using iScript™ Reverse Transcription Supermix for RT-qPCR (Biorad, 1708841). Total expression of transcripts was quantified by qRT-PCR using Powerup SYBR Green Master Mix (Life Technologies, A25778) and amplification performed on a QuantStudio3 (Applied Biosystems). Expression primers can be found in Table S2.

#### Bisulfite conversion and pyrosequencing

To assess methylation, genomic DNA was extracted from pre-plated iPSC’s or sorted RA differentiated populations. The cells were lysed overnight at 50C in Lysis buffer (10mM Tris pH 7.5, 150mM NaCl, 0.2% SDS) followed by isopropanol precipitation. Genomic DNA was sent to EpigenDx Inc. for bisulfite conversion and pyrosequencing. The genomic location (mm10) of the three assessed regions were: Region 1: chr12:109528391-109528471, Region 2: chr12:109530388-109530457 and Region 3: chr12:109541718-109541777.

#### ChIP-qPCR

ChIP was performed as previously described (Liu et al., 2017). Specifically, cells were crosslinked in 1% formaldehyde at room temperature for 10 min and quenched with 125 mM glycine for 5 min at room temperature. The cells were used for KLF4 ChIP (50 × 106 cells) and H3K27ac ChIP (10 × 106 cells). The cell pellets were washed twice in PBS and resuspended in 400 μl lysis buffer (10 mM Tris pH 8, 1 mM EDTA and 0.5% SDS) per 20 × 106 cells. The cells were sonicated using the Bioruptor (Diagenode) (30 cycles of 30 s on/off; high setting) and spun down at the maximum speed for 10 min at 4°C. The supernatants were diluted five times with dilution buffer (0.01% SDS, 1.1% Triton X-100, 1.2 mM EDTA, 16.7 mM Tris pH 8 and 167 mM NaCl) and incubated overnight with antibodies against histone H3K27ac (Abcam, ab4729) with rotation at 4°C. Protein G Dynabeads (Invitrogen, 10004D) pre-blocked with BSA protein (100 ng per 10 μl Dynabeads) were added (10 μl blocked Dynabeads per 10 × 106 cells) the following day and incubated for 2–3h at 4°C. The beads were immobilized on a magnet and washed twice in low-salt buffer (0.1% SDS, 1% Triton X-100, 2 mM EDTA, 150 mM NaCl and 20 mM Tris pH 8), twice in high-salt buffer (0.1% SDS, 1% Triton X-100, 2 mM EDTA, 500 mM NaCl and 20 mM Tris pH 8), twice in LiCl buffer (0.25 M LiCl, 1% NP-40, 1% deoxycholic acid (sodium salt), 1 mM EDTA and 10 mM Tris pH 8) and once in TE buffer. The DNA was then eluted from the beads by incubating with 150 μl elution buffer (1% SDS and 100 mM NaHCO3) for 20 min at 65°C (vortexing every 10 min). The supernatants were collected and reverse crosslinked by incubation overnight at 65°C in the presence of proteinase K. After RNase A treatment for 1 h at 37°C, the DNA was purified using a minElute kit (Qiagen, 28004). Enrichment of protein binding to defined DNA sequences was assessed by qPCR. Primer sequences can be found in supplemental Table S2.

#### 4C-seq

Experiments were performed in duplicate to generate two technical replicates per sample. JF1/B6 iPSC’s (2 × 10^6^) were crosslinked in 1% formaldehyde at room temperature for 10 min and quenched with 125 mM glycine for 5 min at room temperature. The cell pellets were washed twice in PBS and resuspended in 300 μl lysis buffer (10mM Tris-HCl (pH 8.0), 10mM NaCl, 0.2% Igepal CA630 (Sigma I8896), on ice for 20 min. Following centrifugation at 2,500g for 5 min at 4°C, the pellet was resuspended in in 50uL of 0.5% SDS and incubated for 10 min at 65°C. SDS was quenched with 145uL water and 25uL of 10% TritonX-100. 25ul of Cutsmart buffer was added with DpnII enzyme (10 μl; NEB, R0543M) and the samples were incubated overnight at 37 °C with 700rpm rotation. The samples were then diluted with 663μl Milli-Q water, 120 μl T4 ligation buffer (NEB B0202), 60ul ATP 10mM, 120 μl Triton X-100, 12ul BSA 10mg/ml and incubated with Ligase (5 μl, 400U/μl; NEB M0202L) for 3h on a rotor at room temperature. The samples were then treated with proteinase K and reverse crosslinked overnight. Following RNAse treatment, phenol/chloroform extraction and DNA precipitation, the pellets were dissolved in 100 μl of 10 mM Tris pH 7.5 and digested overnight at 37°C by adding 20 μl Cutsmart buffer (NEB), 10ul μl NlaIII (NEB, R0125) and 70 μl Milli-Q water. Following enzyme inactivation, the samples were diluted in 2345 ul Milli-Q water, 300 ul 10×ligation buffer (NEB), 150ul ATP 10mM and incubated with, 5 μl T4 DNA ligase 2M U/ul (NEB, M0202M) overnight at 16°C. The DNA was purified by phenol/chloroform extraction, ethanol precipitation and Zymo columns (D4014).

For allele-specific 4C-seq library preparation, primers were designed upstream of a SNP within IG-DMR. Due to lack of suitable restriction fragments on ΔMat^TRE^ samples, non-allele specific libraries were prepared with the *Gtl2* DMR as viewpoint. Library preparation was further performed by using a PCR strategy as previously described (Krijger et al., 2020). Briefly, 4×200 ng of 4C-template DNA was used to PCR amplify the libraries using the Roche Expand long template PCR system (Roche, 11681842001). Primers were removed using Ampure XP beads (Beckman Coulter, A63880). A second round of PCR was performed using the initial PCR library as a template, with overlapping primers to add the full adapters. The libraries were sequenced on a miSeq platform in SE150 mode. All of the primer sequences can be found in Table S2.

#### Motor neuron differentiation assays

Motor neuron differentiations were performed as described (Wichterle et al., 2002). Briefly, iPSC colonies were dissociated and plated at a 23K cells/cm2 in ADFNK medium (Neurobasal medium (Thermofisher 21103049), 10% KnockOut Serum Replacement (Thermofisher 10828028), 0.1 mM beta-mercaptoethanol, 2mM L-glutamine, and penicillin/streptomycin). Two days post-dissociation, embryoid bodies were treated with 1μM retinoic acid and 0.5μM SAG for four days. Embryoid bodies were collected for further analysis at day 6 post dissociation.

#### Immunocytochemistry

Embryoid bodies were fixed, sectioned, and processed for immunocytochemistry with antibodies as previously described (Novitch et al., 2001). Briefly, embryoid bodies were fixed for 30 minutes at 4°C with 4% Paraformaldehyde, 0.1% Triton-X-100, and 10% Horse Serum in PBS. Then, following 3 washes in 1x PBS, embryoid bodies were equilibrated in 30% sucrose/PBS for 30 minutes, embedded in OCT, and sectioned on a crystostat (15 μM). Primary antibody incubation of sections was performed in PBS with 10% Horse Serum overnight at 4°C. Following 3 washes in 1X PBS, secondary antibody incubation of sections was performed in PBS with 10% Horse Serum for 1 hour at room temperature. Sections were then imaged on an inverted microscope (Zeiss Axio Observer Z1 Inverted Microscope).

### Quantification and statistical analysis

#### Analysis of 4C-seq data

The 4C-seq data was analyzed in a similar fashion as recently described (Raviram et al., 2016, Di Giammartino et al., 2019). Viewpoint primers were trimmed off from all sequencing reads using seqtk (version 1.3.0). Then, allele-specific reads were identified by perfectly matching the respective allele-specific primer sequence against each read, and raw fastq files were split to represent maternal and paternal alleles as individual samples separately. The read-sequence was aligned using bowtie2 v2.3.4.1 (Langmead and Salzberg, 2012) against a reduced genome that consists only of reference genome sequences adjacent to DpnII/NlaIII cut-sites (following the 4C-ker pipeline) (Raviram et al., 2016). Next, the genome was binned into 5kb bins shifted by 500bp (overlapping by 90% with adjacent bins). Reads were counted by unique alignment position in all overlapping bins. Read counts per bin were normalized by sequencing depth per replicate using edgeR (version 3.14.0) (Nikolayeva and Robinson, 2014), resulting in counts per million (CPM). Significance of differential interaction was determined by glmQLFit and glmQLFTest functions from edgeR followed by mutliple testing correction and displaying false-discovery rate (FDR). Visualization was done using average CPM signals per condition.

### Statistical analysis

Results were displayed as mean□±□SEM and statistically analyzed using the GraphPad Prism software (version 8.4.1). Statistical significance of differences between the results was assessed mostly using a two-tailed unpaired Student’s *t*-test. Welch’s correction was applied when comparing samples with significantly different variances. Two-tailed Paired Student’s *t*-test was used specifically for CpG methylation analysis. Statistically significant p-values were indicated as follows: * p□≤□0.05, ** p□≤□0.01, *** p□≤□0.001, **** p□≤□0.0001. No statistical method was used to predetermine sample size, nor blinding or randomization of samples were applied.

## INVENTORY OF SUPPLEMENTAL INFORMATION

Table S1. gRNAs

Table S2. qPCR, ChIP, Surveyor, 4Cseq, Genotyping and Cloning primers

Table S3. Plasmid overview

Table S4. Sample summary

