## Supplemental Material for "A bipartite element with allele-specific functions safeguards DNA methylation imprints at the *Dlk1-Dio3* locus"

**SUPPLEMENTAL INFORMATION : Aronson, Scourzic *et al.***

4 Supplemental Figures (S1 to S4)

**Supplemental Figure 1 (S1).** Related to Figure 1

**Supplemental Figure 2 (S2).** Related to Figure 2

**Supplemental Figure 3 (S3).** Related to Figure 3

**Supplemental Figure 4 (S4).** Related to Figure 4

4 Supplemental Tables

**Table S1:** Guide RNA sequences

**Table S2:** qPCR, ChIP-qPCR, Surveyor, 4C, genotyping and cloning primers

**Table S3:** List of plasmids used in this study

**Table S4:** Description of clones analyzed in this study

Supplemental References

#### SUPPLEMENTAL FIGURES

### Fig S1

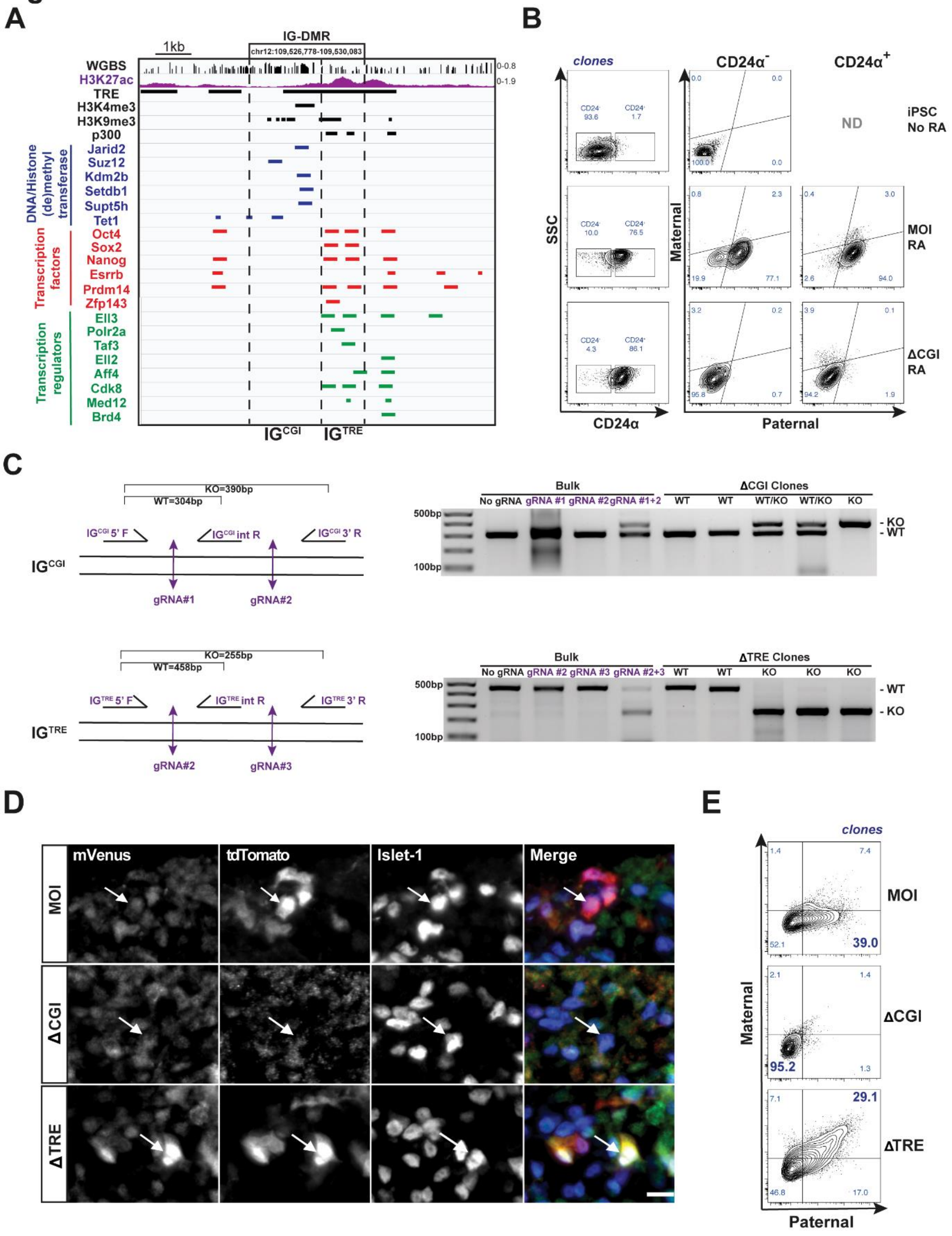

**Figure S1: related to Fig 1**

(**A**) Additional in house (Liu et al., 2017, Di Giammartino et al., 2019) and published ChIPseq datasets (Sanchez-Castillo et al., 2015, Williams et al., 2011) confirm a dichotomy between the $IG^{CGI}$ and $IG^{TRE}$ regarding the occupancy of DNA/histone (de)methyltransferases, transcription factors and regulators. (**B**) Schematic representation of the flow cytometry strategy used to assess iPSC and RA-differentiated cells using a $CD24a^{+}$ gating and subsequent *Dlk1* reporter expression (mVenus and tdTomato). This strategy ensures that Dlk1-loss clones are not inefficiently differentiated, but indeed repress the *Dlk1* gene itself. ND= Not done. (**C**) Genotyping strategy used to confirm $IG^{CGI}$ and $IG^{TRE}$ deletions (ΔCGI and ΔTRE) using 2 gRNAs (left). Complete list of gRNAs and primers used are provided in Tables S1 and S2. Representative agarose gel showing specific detection of wild type (WT) and knockout (KO) bands (right). (**D**) Immunofluorescence detection of Venus and tdTomato expression as well as the neuronal marker Islet-1 demonstrate bi-allelic expression in a representative $IG^{TRE}$ clone. Neither mVenus nor tdTomato is detected for Islet-1 stained cells in $IG^{CGI}$ deleted clones. Bar= 40μM (**E**) Representative FACS plots of WT, ΔCGI and ΔTRE clones after directed differentiation to motor neurons confirm the MOI, LOI-Dlk1loss and LOI-BiDlk1 phenotypes.

Fig S2

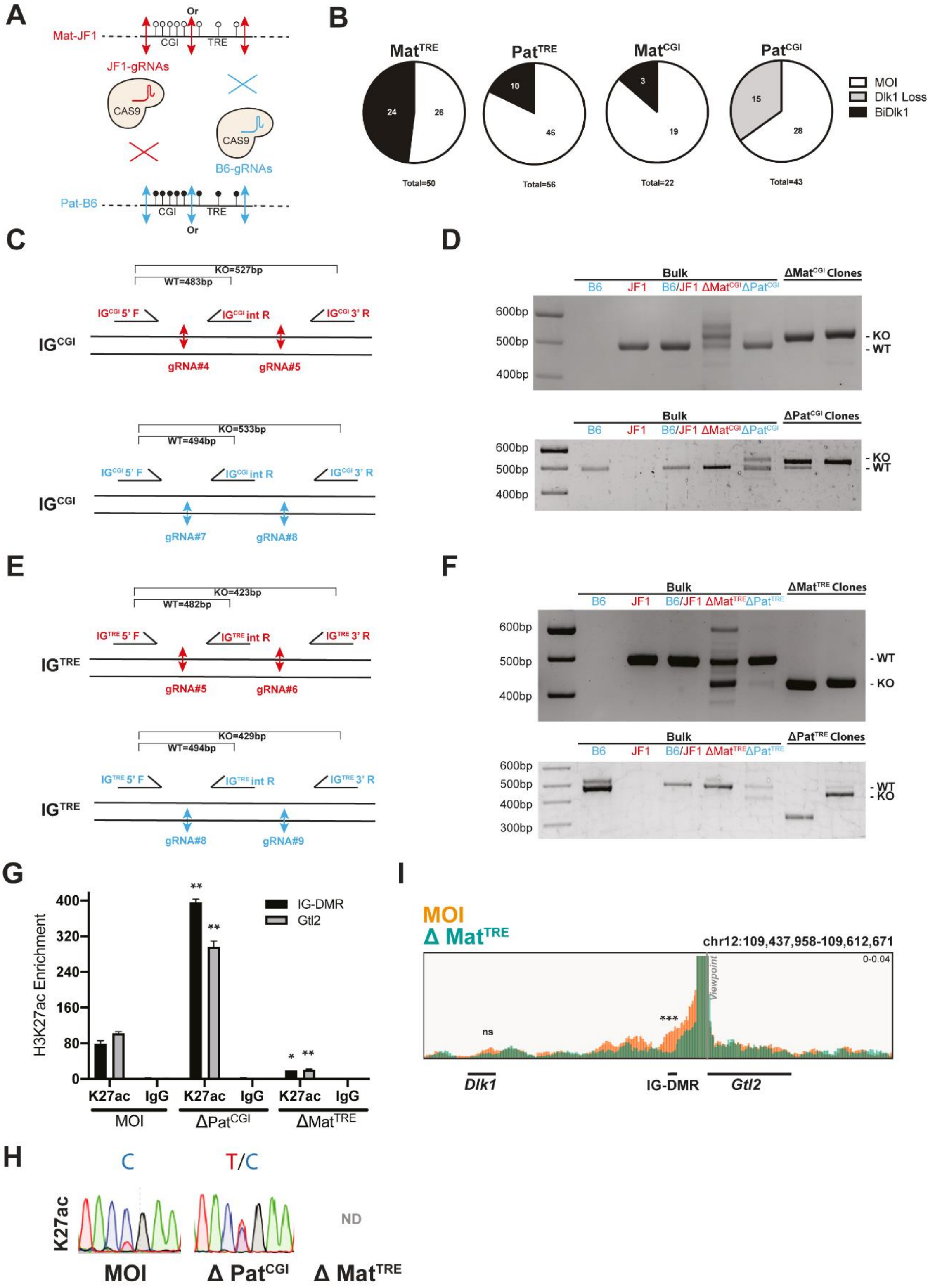

A
Mat-JF1
Or
CGI
TRE
JF1-gRNAs
CAS9
B6-gRNAs
Pat-B6
B
MatTRE
PatTRE
MatCGI
PatCGI
Total=50
Total=56
Total=22
Total=43
MOI
Dlk1 Loss
BiDlk1
C
KO=527bp
WT=483bp
IGCGI 5' F
IGCGI int R
IGCGI 3' R
gRNA#4
gRNA#5
KO=533bp
WT=494bp
gRNA#7
gRNA#8
D
Bulk
ΔMatCGI Clones
ΔPatCGI Clones
B6
JF1
B6/JF1
ΔMatCGI
ΔPatCGI
600bp
500bp
400bp
- KO
- WT
E
KO=423bp
WT=482bp
IGTRE 5' F
IGTRE int R
IGTRE 3' R
gRNA#5
gRNA#6
KO=429bp
WT=494bp
gRNA#8
gRNA#9
F
ΔMatTRE Clones
ΔPatTRE Clones
ΔMatTRE
ΔPatTRE
300bp
G
H3K27ac Enrichment
IG-DMR
Gtl2
K27ac
IgG
MOI
ΔPatCGI
ΔMatTRE
H
C
T/C
K27ac
ND
Δ PatCGI
Δ MatTRE
I
MOI
Δ MatTRE
chr12:109,437,958-109,612,671
0-0.04
Viewpoint
ns
***
Dlk1
IG-DMR
Gtl2

**Figure S2: related to Fig 2**

(**A**) Schematic representation of CRISPR/Cas9 targeting using allele-specific gRNAs, corresponding to SNPs identified between JF1/B6 murine backgrounds. (**B**) Pie charts showing the distribution of phenotypes based on FACS analysis of multiple clones isolated after allele-specific targeting of $IG^{CGI}$ and $IG^{Enh}$. Numbers of clones are shown the charts. (**C**) Strategy used for allele-specific genotyping of $IG^{CGI}$ targeted clones and (**D**) representative agarose gel showing the distinct wild type (WT) and knockout (KO) DNA bands. (**E**) Strategy used for allele-specific genotyping of $IG^{TRE}$ clones and (**F**) Representative agarose gels showing the respective WT and KO bands. (**G)** H3K27ac enrichment at IG-DMR (downstream of the $IG^{TRE}$) and *Gtl2* promoter assessed by ChIP-qPCR in N=2 clones from each indicated genotype. Statistical significance relative to their respective MOI clones was calculated by two-tailed paired Student's t-test. (**H)** Allele-specific sequencing of H3K27ac ChIP-qPCR product in a $\Delta Pat^{CGI}$ clone show a biallelic enrichment (C/T) in contrast with a normally imprinted clone which detects only the maternal allele (C). (**I**) Non allele-specific 4C-seq, using the *Gtl2* promoter as viewpoint, demonstrates interactions with the IG-DMR and *Dlk1* in MOI iPSCs and reduced interactions upon $Mat^{TRE}$ deletion. The IG-DMR lacks suitable frag-ends in $\Delta Mat^{TRE}$ clones and could therefore not be chosen as viewpoint. Statistics for *Gtl2* interaction with the IG-DMR and *Dlk1* are shown at chr12: 109526369-109529934 (FDR= 0,0045) and chr12:109453455-109463336 (FDR= 0,999) respectively. *Asterisks indicate significance: ** p<0.01. Error bars represent +/- SEM.*

Fig S3

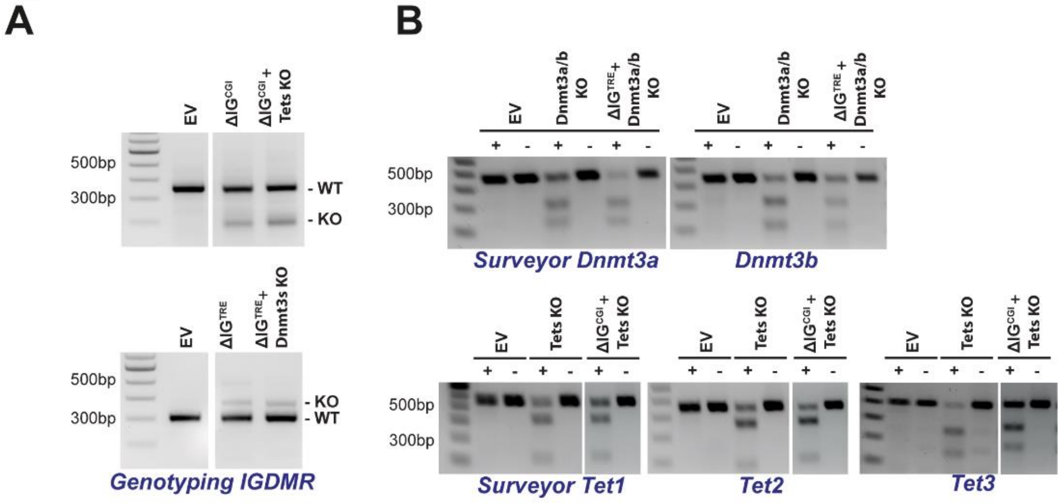


**<u>Figure S3: related to Fig 3</u>**

(**A)** Examples of genotyping agarose gels that show the detection of wild type (WT) and knockout (KO) bands upon targeting of either IG-CGI (top) or IG-TRE (bottom). (**B**) Examples of agarose gels showing the results of surveyor assays that used to detect indels as a measure of successful CRISPR/Cas9-targeting of Dnmt3a and Dnmt3b (top) and Tet1, Tet2 and Tet3. Importantly, similar levels of targeting were observed across conditions.

#### Fig S4

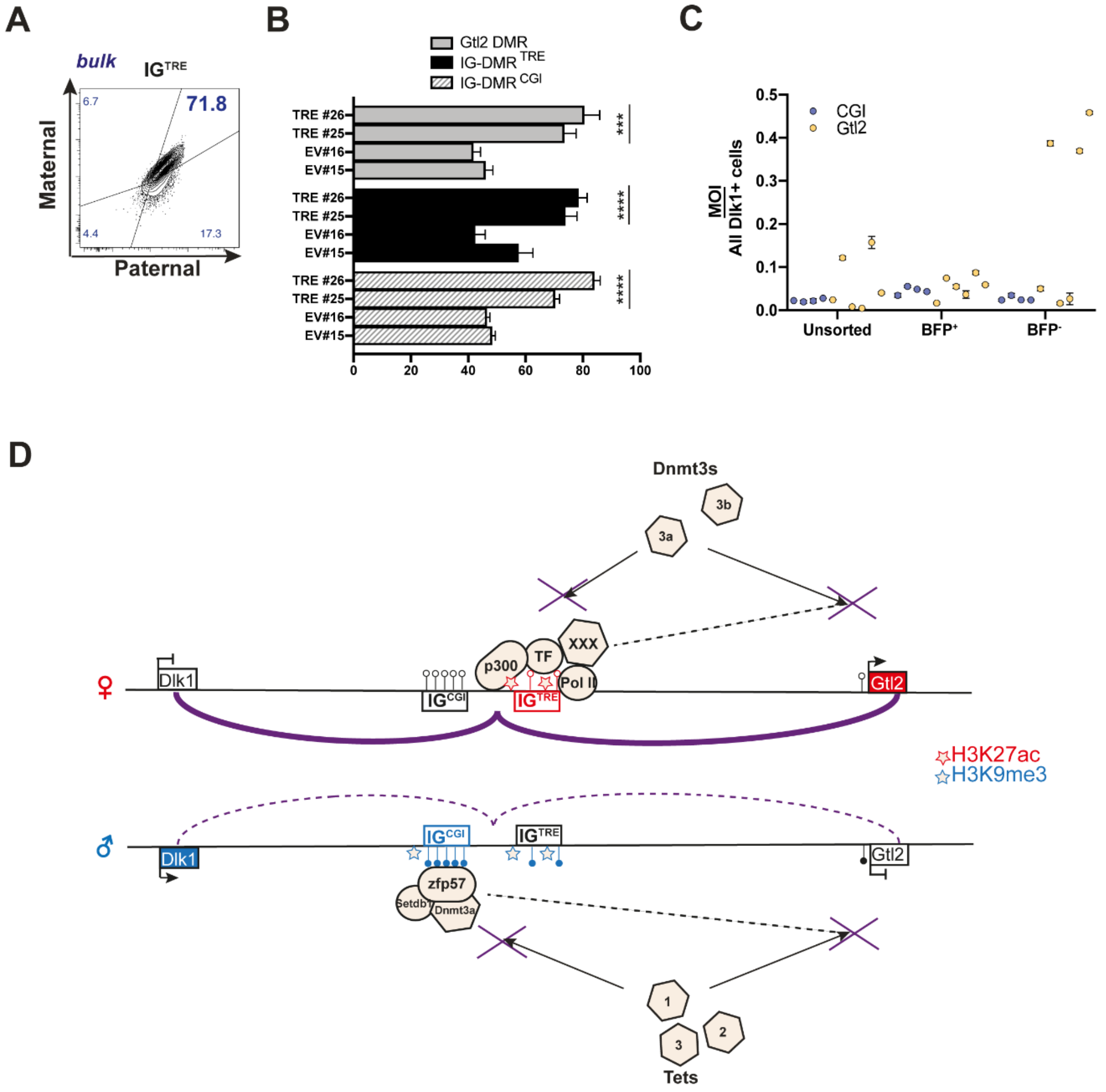


**Figure S4: related to Fig 4**

(**A**) Representative FACS plot and (**B**) DNA methylation levels of IG$^{TRE}$-targeted clones with dCas9-BRF-KRAB. Statistical analysis of each DMR (and each clone) is compared to the EV #15 and 16 using two-tailed unpaired t-test. Means are represented as +/- SEM. (**C**) FACS quantitation showing the percentage of MOI populations of N=4 dCas9-BFP-KRAB CGI-targeted clones and N=6 Gtl2-targeted clones at p15. Means are represented as +/- SEM. (**D**) Model figure of the proposed mechanisms that ensure MOI at *Dlk1-Dio3*. Zfp57, Dnmt3s and Setdb1 factors have been shown to be recruited to the paternally inherited methylated IG$^{CGI}$ (blue). In contrast, p300 and transcriptional machinery together with cofactors are associated with the maternally inherited unmethylated IG$^{TRE}$ (red). Crossed arrows represent the Tets and Dnmt3s activity prevention mediated by IG$^{CGI}$ and IG$^{TRE}$, respectively. The protective effects of

IG-DMR protection on the methylation and expression of *Gtl2* is represented with dashed arrows. Thick plain lines represent strong long-range chromatin contacts on the maternal allele, while thin dashed lines represent weaker interactions on the paternal allele. Stars indicates H3K27ac mark deposition on the maternal allele (red) or H3K9me3 on the paternal allele (blue).

#### SUPPLEMENTAL TABLES

##### Table S1: gRNA's

| gRNA | Name | Sequence oligo | Reference |
|---|---|---|---|
| #1 | 5' $IG^{CGI}$ | CACCGTGACACAACAGTCTTGACC<br>AAACGGTCAAGACTGTTGTGTCAC | This paper |
| #2 | 3' $IG^{CG}$I<br>/5' $IG^{TRE}$ | CACCGGAGACAAGAACTCCGAGAC<br>AAACGTCTCGGAGTTCTTGTCTCC | |
| #3 | 3' $IG^{TRE}$ | CACCGATATCTCTCACCTGACTAA<br>AAACTTAGTCAGGTGAGAGATATC | |
| #4 | mat-5' $IG^{CGI}$ | CACCGCTCAGCACAACTACATGCA<br>AAACTGCATGTAGTTGTGCTGAGC | |
| #5 | mat-3' $IG^{CGI}$<br>/mat-5' $IG^{TRE}$ | CACCGTCACAACGCCTCCAGCTAA<br>AAACTTAGCTGGAGGCGTTGTGAC | |
| #6 | mat-3' $IG^{TRE}$ | CACCGAACCTCTGATCAGCAGGCC<br>AAACGGCCTGCTGATCAGAGGTTC | |
| #7 | pat-5' $IG^{CGI}$ | CACCGCCACCTCAGCACAACTACA<br>AAACTGTAGTTGTGCTGAGGTGGC | |
| #8 | pat-3' $IG^{CGI}$<br>/pat-5' $IG^{TRE}$ | CACCGACAGTTCCCAGGCAGCCCT<br>AAACAGGGCTGCCTGGGAACTGTC | |
| #9 | pat-3' $IG^{TRE}$ | CACCGAACCTCTGATCAGCAGGCT<br>AAACAGCCTGCTGATCAGAGGTTC | |
| #10 | Tet1 | CACCGGCTGCTGTCAGGGAGCTCA<br>AAACTGAGCTCCCTGACAGCAGCC | (Wang et al., 2013) |
| #11 | Tet2 | CACCGAAAGTGCCAACAGATATCC<br>AAACGGATATCTGTTGGCACTTTC | |
| #12 | Tet3 | CACCGAAGGAGGGGAAGAGTTCTCG<br>AAACCGAGAACTCTTCCCCTCCTTC | |
| #13 | Dnmt3a ex 17 | CACCGTGGGCATGGTGCGGCACCA<br>AAACTGGTGCCGCACCATGCCCAC | (Sano et al., 2018) |
| #14 | Dnmt3b ex 19 | CACCGGAGTGGGGCCCGTTCGACT<br>AAACAGTCGAACGGGCCCCACTCC | (Dai et al., 2016) |
| #15 | CpG site -1 | CACCGATTGTGCCGCGGTTCGCCG<br>AAACCGGCGAACCGCGGCACAATC | This paper |
| #16 | CpG site -2 | CACCGTCGCGGCACGCGTACACAG<br>AAACCTGTGTACGCGTGCCGCGAC | |
| #17 | CpG site -3 | CACCGCTACCGCTACGGTTCATAG<br>AAACCTATGAACCGTAGCGGTAGC | |
| #18 | Proseq site -1 | CACCGAAATCTGGCACTCCGTTTC<br>AAACGAAACGGAGTGCCAGATTTC | |
| #19 | Proseq site -2 | CACCGTGAGCAGAGACGAGTTGGC<br>AAACGCCAACTCGTCTCTGCTCAC | |
| #20 | Proseq site -3 | CACCGTATTCTAGGGTTTGTCACC<br>AAACGGTGACAAACCCTAGAATAC | |
| #21 | Proseq site -4 | CACCGCAGTGTGCCAGTTTTTTCG<br>AAACCGAAAAAACTGGCACACTGC | |
| #22 | Gtl2 prom -1 | CACCGCGTCTTCTGTGCTAGGGGC<br>AAACGCCCCTAGCACAGAAGACGC | |
| #23 | Gtl2 prom -2 | CACCGTCCTCCTGGACATGCCGAA<br>AAACTTCGGCATGTCCAGGAGGAC | |
| #24 | Gtl2 prom -3 | CACCGGGATGGCTAACCACTCACC<br>AAACGGTGAGTGGTTAGCCATCCC | |

**Table S2: qPCR, ChIP-qPCR, Surveyor, 4C, genotyping and cloning primers**

| Gene/region | Sequence | Reference |
|---|---|---|
| **qPCR primers** | | |
| qPCR_Gtl2 cds F | TTGCACATTTCCTGTGGGAC | (Stadtfeld et al., 2010) |
| qPCR_Gtl2 cds R | AAGCACCATGAGCCACTAGG | |
| qPCR_Dlk1 cds F | CCCAGGTGAGCTTCGAGTG | |
| qPCR_Dlk1 cds R | GGAGAGGGGTACTCTTGTTGAG | |
| qPCR_Gapdh cds F | AGGTCGGTGTGAACGGATTTG | |
| qPCR_Gapdh cds R | TGTAGACCATGTAGTTGAGGTCA | |
| **ChIP-qPCR primers** | | |
| ChIP_IG$^{TRE}$ DS F | TGATTCACCACAATTCGTTCTGG | This paper |
| ChIP_IG$^{TRE}$ DS R | CCTGGCGAGCAACAGTGAAGCTT | |
| ChIP_Gtl2 DMR F | GGTAGAGACCCTGATTAAGAAAAG | (Stadtfeld et al., 2010) |
| ChIP_Gtl2 DMR R | CTCTCACCATTAATGCGATTC | |
| **Surveyor primers** | | |
| Surv_Tet1 F | CCTCAACACAAGTCTCGGAAGAC | This paper |
| Surv_Tet1 R | CTGTGCTATTTGTAAAACTGGGTC | |
| Surv_Tet2 F | CTGAAACATCGTCGAGAACAGAAC | |
| Surv_Tet2 R | CAATCCCAGCCTGGTCATAAC | |
| Surv_Tet3 F | GCTGCCACAAGGGTCATTG | |
| Surv_Tet3 R | CAGAGGAGGAGCATCTGGAAATG | |
| Surv_Dnmt3a ex 17 F | CTTGATTTGGGAAAGGATACTGGG | |
| Surv_Dnmt3a ex 17 R | CAAATGGGCTGAATGGAAGGAG | |
| Surv_Dnmt3b ex 19 F | GCAAACAGGGCAGTTGGAG | |
| Surv_Dnmt3b ex 19 R | GAACATAGACACATGGCACATAGTGG | |
| **4C primers** | | |
| 4C_PCR1-IG$^{TRE}$-NOI1 R | TACACGACGCTCTTCCGATCTGGGAAAAGT CAAAATGGTCACCCTAACCC | This paper |
| 4C_PCR1-IG$^{TRE}$-NOI2 R | TACACGACGCTCTTCCGATCTCTGGGAAAA GTCAAAATGGTCACCCTAACCC | |
| 4C_PCR1-IG$^{TRE}$-Dlk1Loss-1 R | TACACGACGCTCTTCCGATCTTTAAGGGGG AAAAGTCAAAATGGTCACCCTAACCC | |
| 4C_PCR1-IG$^{TRE}$-Dlk1Loss-2 R | TACACGACGCTCTTCCGATCTGACTGGGAA AAGTCAAAATGGTCACCCTAACCC | |
| 4C_PCR1-IG$^{TRE}$ F | ACTGGAGTTCAGACGTGTGCTCTTCCGATC TACCCCCCAGTAGCTTGTAAGCCATG | |
| 4C_PCR1-Gtl2-DMR-NOI1 R | TACACGACGCTCTTCCGATCTTCAGCCCAA GCAGACACATG | |
| 4C_PCR1-Gtl2-DMR-NOI2 R | TACACGACGCTCTTCCGATCTCTCAGCCCA AGCAGACACATG | |
| 4C_PCR1-Gtl2-DMR-BiDlk1-1 R | TACACGACGCTCTTCCGATCTACTCAGCCC AAGCAGACACATG | |
| 4C_PCR1-Gtl2-DMR-BiDlk1-2 R | TACACGACGCTCTTCCGATCTGACTCAGCC CAAGCAGACACATG | |
| 4C_PCR1-Gtl2-DMR F | ACTGGAGTTCAGACGTGTGCTCTTCCGATC TCTGTCTACAAATCGCCCTTCCATC | |
| 4C_PCR2-index2 F | CAAGCAGAAGACGGCATACGAGATCCGTAG GTGACTGGAGTTCAGACGTGTGCT | |
| 4C_PCR2-index17 F | CAAGCAGAAGACGGCATACGAGATTCTACA GTGACTGGAGTTCAGACGTGTGCT | |
| 4C_PCR2-index64 F | CAAGCAGAAGACGGCATACGAGATTTAAGG GTGACTGGAGTTCAGACGTGTGCT | |

| | | |
|---|---|---|
| 4C_PCR2-index67 F | CAAGCAGAAGACGGCATACGAGATCGGTGT GTGACTGGAGTTCAGACGTGTGCT | This paper |
| 4C_PCR2-index75 F | CAAGCAGAAGACGGCATACGAGATGAACGG GTGACTGGAGTTCAGACGTGTGCT | |
| 4C_PCR2-index82 F | CAAGCAGAAGACGGCATACGAGATGGCGTA GTGACTGGAGTTCAGACGTGTGCT | |
| 4C_PCR2 R | AATGATACGGCGACCACCGAGATCTACACT CTTTCCCTACACGACGCTCTTCCGATCT | |
| **Genotyping primers** | | |
| Geno_IG$^{CGI}$ 5' flank | GCTCAGGTTCAAGTCGGCTAC | This paper |
| Geno_IG$^{CGI}$ 3' flank | GATGAGACTCCTCTAGGATTCCC | |
| Geno_IG$^{CGI}$ inside | CCTGTTCCCTAGTGAAGGTTTGC | |
| Geno_IG$^{TRE}$ 5' flank | CACAGTTCCCAGGCAGCCCTTG | |
| Geno_IG$^{TRE}$ 3' flank | CACTAGCTGGTATTGTATAAAACC | |
| Geno_IG$^{TRE}$ inside | GGAGCCTTGAGCCCAGAAC | |
| Geno_Mat$^{CGI}$ 5' flank | GTGGGGTTGGTTGCCTAAT | |
| Geno_Mat$^{CGI}$ 3' flank | GATGAGACTCCTCTAGGATTCCTG | |
| Geno_Mat$^{CGI}$ inside | GGAATCTGGTACTTGCATAGTTC | |
| Geno_Pat$^{CGI}$ 5' flank | GTGGGGTTGGTTGCCGAGC | |
| Geno_Pat$^{CGI}$ 3' flank | GATGAGACTCCTCTAGGATTCACA | |
| Geno_Pat$^{CGI}$ inside | GGAATCTGGTACTTGCAGAATTT | |
| Geno_Mat$^{TRE}$ 5' flank | GGAATCTGGTACTTGCATAGTTC | |
| Geno_Mat$^{TRE}$ 3' flank | CTAGTCTTGGGAGTGAAATTAC | |
| Geno_Mat$^{TRE}$ inside | GATGAGACTCCTCTAGGATTCCTG | |
| Geno_Pat$^{TRE}$ 5' flank | GGAATCTGGTACTTGCAGAATTT | |
| Geno_Pat$^{TRE}$ 3' flank | CTAGTCTTGGGAGTGAAACGAT | |
| Geno_Pat$^{TRE}$ inside | GATGAGACTCCTCTAGGATTCACA | |
| **Cloning primers** | | |
| EF1a-promoter F | CCGGAATTCCGGACTAGTCGTGAG GCTCCG | This paper |
| EF1a-promoter R | CCGGAATTCCGGGCTTCACGACAC CTGAAATGG | |
| dCas9-Dnmt3a (Gibson) F | CAGGTGTCGTGAAGCCCGGAATTCTC ACATGGACTATAAGGACCACGAC | |
| dCas9-Dnmt3a (Gibson) R | CAGTAACGTTAGGGGGGGGGGGCGG TCACACACACGCAAAATACTCCTTCAG | |
| dCas9-tet1 (Gibson) F | CAGGTGTCGTGAAGCCCGGAATTCTC ACATGGACTATAAGGACCACGACG | |
| dCas9-tet1 (Gibson) R | CAGTAACGTTAGGGGGGGGGGGCGG CTCAGACCCAATGGTTATAGGGCC | |
| IRES-BSD F | CCCTTGAACCTCCTCGTTCGA | |
| IRES-BSD (Gibson) R | CGATAAGCTTGATATCAAGCTTGCATGC CTGCATTAGCCCTCCCACACATAACCAG | |
| U6-guide F (XbaI) | ATGCTTCTAGAGAGGGCCTATTTCCCATGATT | (Dow et al., 2015) |
| U6-guide F (EcoRI) | ATGCTGAATTCGAGGGCCTATTTCCCATGATT | |
| U6-guide F (BamHI) | ATGCTGGATCCGAGGGCCTATTTCCCATGATT | |
| U6-guide F (XbaI) | ATGCTCTCGAGGAGGGCCTATTTCCCATGATT | |
| U6-guide R (MfeI) | TGACACAATTGCAAAAAAGCACCGACTCG | |
| U6-guide R (BglII) | TGACAAGATCTCAAAAAAGCACCGACTCG | |
| U6-guide R (SalI) | TGACAGTCGACCAAAAAAGCACCGACTCG | |
| U6-guide R (AvrII) | TGACACCTAGGCAAAAAAGCACCGACTCG | |

**Table S3: Plasmids overview**

| Plasmid | Identifier | Reference |
|---|---|---|
| pdCas9-Dnmt3a-puro | Add Gene #71667 | (Vojta et al., 2016) |
| pPGKENTRY-dCas9-Tet1CD | N/A | (Verma et al., 2018) |
| pMIGR1-IRES-BSD | | N/A |
| pBS-Dlk1-tomato | | (Swanzey and Stadtfeld, 2016) |
| pHR-SFFV-dCAS9-BFP-KRAB | Add Gene #46911 | (Gilbert et al., 2013) |
| pLKO5.sgRNA.EFS.PAC | Add Gene #57825 | (Heckl et al., 2014) |
| pSpCas9(BB)-2A-GFP (px458) | Add Gene #48138 | (Ran et al., 2013) |
| pHR-EF1a-dCAS9-BFP-KRAB | N/A | This paper |
| pHR-EF1a-dCAS9-Dnmt3a-BSD | | |
| pHR-EF1a-dCAS9-Tet1-BSD | | |
| pLKO5.sgRNA.EFS.PAC- Neo | | |

**Table S4: Samples overview**

| Samples | Targeted region | Bulk/Clone |
|---|---|---|
| #1 | $IG^{CGI}$ deletion | clone |
| #2 | | clone |
| #3 | $IG^{TRE}$ deletion | clone |
| #4 | | clone |
| #5 | $Pat^{CGI}$ deletion | clone |
| #6 | | clone |
| #7 | $Mat^{TRE}$ deletion | clone |
| #8 | | clone |
| #9 | dCas9-Tet1 no gRNA | Bulk |
| #10 | dCas9-Tet1 on $IG^{CGI}$ | clone |
| #11 | | clone |
| #12 | dCas9-Dnmt3a no gRNA | Bulk |
| #13 | dCas9-Dnmt3a on $IG^{TRE}$ | clone |
| #14 | | clone |
| #15 | dCas9-KRAB no gRNA | Bulk |
| #16 | | Bulk |
| #17 | dCas9-KRAB on Gtl2 | Bulk |
| #18 | | Bulk |
| #19 | dCas9-KRAB on $IG^{CGI}$ | Bulk |
| #20 | | Bulk |
| #21 | dCas9-KRAB on $IG^{CGI}$ $BFP^{-}$ | clone |
| #22 | | clone |
| #23 | dCas9-KRAB on Gtl2 rescue | clone |
| #24 | | clone |
| #25 | dCas9-KRAB on $IG^{TRE}$ | Bulk |
| #26 | | Bulk |
